# SPaRTA: Spatial Reasoning in Text-to-SQL for Omics Analysis

**DOI:** 10.64898/2026.10.31.763119

**Authors:** Arka Dutta, Souvik Ghosh, Kenneth Prado, Ashiqur R. KhudaBukhsh, Hirak Sarkar

## Abstract

Spatial-omics technologies provide clinicians, pathologists, and biomedical researchers with high-resolution maps of transcriptomic, proteomic, and metabolomic activity within tissues. However, querying these complex datasets often requires specialized computational expertise, limiting their accessibility to domain experts without programming experience. In the current manuscript, we develop `SPaRTA` (Spatial Reasoning in Text-to-SQL for Omics Analysis), a natural-language interface that translates free-text questions into executable spatial `SQL` queries over such data. We release `SPaRTA`-Bench, comprising dataset-specific database schemas, explicit query conventions, and 7,500 question–SQL pairs validated through LLM-assisted verification and targeted human review, spanning breast cancer, pancreatic cancer, and melanoma tissue sections, and identify 15 failure modes in which a query runs but returns a wrong answer. Openweight large language models (LLMs) of the `Llama`, `Mistral`, `Qwen`, `Gemma` and `Phi` families reach 10–40% execution accuracy zero-shot, and at most 61% with reasoning and execution feedback. Fine-tuning raises them to 63–77%, and they lose at most 6 percentage points on a tissue unseen in training. Execution feedback repairs queries that fail to run, but rarely queries that run and return a wrong answer.

**Code and data:** github.com/sarkar-lab/SPARTA_public

## 1 Introduction

Advances in imaging and sequencing technologies (Ståhl et al., 2016; de Oliveira et al., 2025; 10x Genomics, 2026; Rodriques et al., 2019; Stickels et al., 2021) now profile biological tissue at high resolution and scale. Depending on the platform, these spatial-omics datasets (where *omics* stands for modalities such as RNA and protein) measure thousands of genes, proteins, metabolites and other modalities, across millions of cells, along their spatial context (Song et al., 2025). These datasets reveal how cells interact (e.g., tumors interacting with immune cells) or which tissue regions respond to a specific treatment. However, a single spatialomics dataset can contain millions of cells, each characterized by hundreds or thousands of molecular measurements, together with their spatial coordinates and tissue context. Answering biological questions requires navigating these measurements and their spatial relationships, often through complex computational and geometric analyses. This makes such datasets particularly difficult to explore for clinicians, pathologists, and biologists without computational training.

Researchers increasingly use `LLM` chatbots such as `ChatGPT`, `Claude` to work with clinical data (Thirunavukarasu et al., 2023), but two problems make them unsuitable for spatial-omics data. First, clinical datasets are sensitive and large. Uploading them to an external service is impractical and raises privacy and security concerns (Faustini et al., 2026; Safarlou et al., 2021), including compliance with HIPAA (Marks and Haupt, 2023). Results from a proprietary model are also hard to reproduce, since its API can change without notice (Chen et al., 2024) and the model can be retired (Wolfrath et al., 2026). Second, generative models hallucinate (Huang et al., 2025; Bang et al., 2025; Ravichander et al., 2025), and in clinical applications hallucination can be harmful. A hallucinated threshold, for example, can mark the cells of a patient sample as cancerous.

An open-weight `LLM` deployed locally addresses both challenges by translating natural-language questions into structured query language (`SQL`) queries, a task known as natural-language-to-SQL (NL2SQL). Given a database schema and a natural-language question, the model generates an SQL query that can be executed directly against the dataset. NL2SQL has advanced considerably with `LLM`-based methods (Liu et al., 2025; Talaei et al., 2024; Pourreza and Rafiei, 2024; Li et al., 2024), whose performance is increasingly evaluated through challenging benchmarks (Li et al., 2023; Lei et al., 2025). While domain-specific benchmarks exist for clinical records (Wang et al., 2020; Lee et al., 2022), scientific databases (Zhang et al., 2024), biomedical knowledge bases (Koretsky et al., 2026), and geographic queries (Hou et al., 2025; Kazazi et al., 2025), none specifically evaluates the combination of molecular, cellular, and spatial reasoning needed to query spatial-omics datasets. A related benchmark SpatialBench (Workman et al., 2025) addresses spatial biology questions but focuses on generating analysis code rather than SQL queries.

General `SQL` benchmarks also do not reflect the kinds of questions clinicians and pathologists ask about tissue. For example, a pathologist examining breast cancer spatial-omics data may ask:

`“Which spatial domains show the highest fraction of cytotoxic T cells expressing GZMB that are proximal to proliferating tumor cell clusters?”`.

A general benchmark asks instead:

`“Find the names of students who are enrolled in more than 3 courses and have a GPA above 3.5”` Only the first question needs implicit biological knowledge and spatial predicates.

### 1.1 Contributions

In this paper, we present `SPaRTA`: **Spa**tial **R**easoning in **T**ext-to-SQL for Omics **A**nalysis. Drawing on domain knowledge (Bogdanov et al., 2026) and iterative improvement (Liu et al., 2022; Shi et al., 2025), we make three contributions:

#### A benchmark for spatial-omics NL2SQL

We release `SPaRTA`-Bench : a database schema extracted from three real Xenium datasets (breast cancer, pancreatic cancer and melanoma), written conventions, and 7, 500 question–`SQL` pairs from 15 query categories, spanning spatial proximity to multi-hop reasoning over expression, neighborhood composition and tissue architecture (Figure 1 a-e). An agent-based pipeline generates the pairs, independent re-derivation and blind audits verify them, and annotators with domain knowledge validate a sample (Figure 1 d). Template-held-out and tissue-heldout splits share no template between training and test.

**Figure 1.**
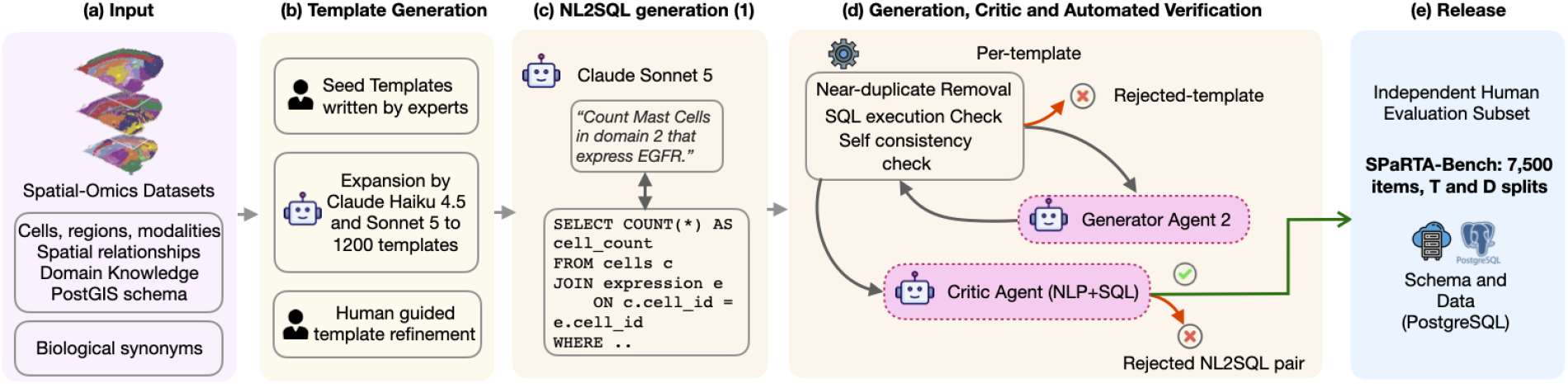
The benchmark construction pipeline of `SPaRTA` (Section 3). (a) Spatial-omics datasets, database schemas, and biological conventions serve as inputs. (b) Domain experts and LLMs construct and refine query templates. (c) Templates are instantiated into natural-language questions and corresponding SQL queries. (d) Candidate pairs undergo automated verification, with rejected pairs iteratively revised by generator and critic agents. (e) Independent audits and human validation yield 7,500 question–SQL pairs, organized into template-held-out (T) and dataset-held-out (D) splits.

#### How NL2SQL models fail in biological domains

Off-the-shelf `LLM`s often write plausible `SQL` that runs without error but returns a biologically wrong answer. We identify 15 such failure modes in three areas (structural, faithfulness and semantic errors) and measure how often six open-weight models make them. Execution feedback repairs queries that fail to run, but rarely the ones that return a wrong answer.

#### Fine-tuning open-weight **LLM**s

Fine-tuning five open-weight models on `SPaRTA`-Bench with lowrank adaptation (LoRA; Hu et al., 2022) raises execution accuracy by 35–58 points, and the gains transfer to a tissue unseen in training. Preference optimization does not add to these gains.

## 2 Design Considerations

Spatial-omics questions differ from those of general benchmarks such as `SPIDER` (Yu et al., 2018; Lei et al., 2025) in two ways, which shaped the design of `SPaRTA`.

### Implicit biological concepts

When a pathologist asks about *infiltrating immune cells near a DCIS lesion*, the question presupposes knowledge of cell type hierarchies, marker-cluster correspondences and tissue architecture that lies outside the database schema. Before writing any `SQL`, a model must decide which cell types count as *immune cells*, which clusters represent *DCIS* (there may be several, such as *DCIS_1* and *DCIS_2*) and what radius counts as *near*. Terms also carry technical meanings. In everyday language, *infiltration* means movement or presence. In spatial pathology it means a density, the number of cytotoxic lymphocytes within a set radius of a tumor cell, which a query must compute as a neighborhood count rather than a proximity check. Sharing the schema is therefore not enough, and `SPaRTA` fixes these choices for each dataset in a written conventions document (Section 3).

### Correctness of queries

A query can run, return plausible numbers and still be wrong. Consider a question from `SPaRTA`-Bench, *“compare average GZMB expression of CD8+ T cells within 50 micrometers of invasive tumor cells versus those that are not*.*”* A generated query averages `e.raw_count` over an inner join with the expression table and returns two numbers, one for proximal and one for distal cells. But expression is stored sparsely, and an absent row means a zero count, so the inner join drops every *CD8+ T* cell without a *GZMB* entry and inflates both averages. If proximal cells more often have zero expression, the proximal average is inflated more, and the query can **reverse the conclusion about immune activation near the tumor**. We call such errors *silent failures*: execution alone does not reveal them, yet they change the scientific conclusion.

#### 2.1 Evaluated Models

We consider six open-weight models that run on a single 48 GB GPU: Qwen3-8B (Yang et al., 2025), Qwen3.5-9B (Qwen Team, 2026), Llama 3.1 8BInstruct (Grattafiori et al., 2024), Gemma-4-12Bit (Team et al., 2026), Phi-4 (Abdin et al., 2024), and Mistral-7B-Instruct-v0.3 (Jiang et al., 2023) (Section C). Qwen3, Qwen3.5 and Gemma 4 can reason before they answer. We fine-tune all of them except Qwen3.5-9B, whose linear-attention layers our trainer does not support (Appendix D). As a reference, we also score GPT-6.1-sol, `Claude` Sonnet 5.5 and Gemini 3.8 Flash zero-shot in direct mode, with three runs at each API’s low reasoning setting (Section C). Earlier `Claude` and GPT models helped build the gold (Section 3), which may favor these two families (Xu et al., 2025).

## 3 Dataset and Benchmark Construction

We built the benchmark over three Xenium tissue datasets in five phases. Its queries cover 15 categories and four levels each of `SQL` and spatial complexity.

### Phase 1 Few-shot foundation

We used the 3, 744 syntax-level tasks from GeoSQL-Bench (Hou et al., 2025), each with an explicit and an underspecified question (7,488 NL2SQL pairs), as a reference for `PostGIS` syntax. Generation uses a subset F of `PostGIS` functions for proximity, containment, region properties, spatial aggregation, derived regions, tessellation and shape comparison (Section F).

### Phase 2 Schema extraction

We loaded three Xenium spatial transcriptomics sections, breast cancer (Janesick et al., 2023), pancreatic cancer (10x Genomics, 2023b) and melanoma (10x Genomics, 2023a), from `h5ad` files into one `PostGIS` database each. Cell coordinates, cluster labels *C*, the sparse expression matrix 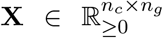 (*n*_*c*_ cells, *n*_*g*_ genes), domain labels *D* and region geometries fill four tables (`cells`, `expression`, `regions`, `genes`) under one shared schema *S* (Table 1 gives the sizes). The `expression` table holds one row per gene detected in a cell, so a missing row means a zero count. The *entity vocabulary* (referred to as the vocabulary) *V* = (*C, G, D*) of cluster labels, gene names *G* and domain labels comes from the same files, so templates use only entities present in the data (Figure 1 a).

**Table 1.** `SPaRTA`-Bench by dataset. Templates: those of the 15 released categories that run on the data (for pancreas and melanoma, ported from breast cancer). Question similarity is the mean pairwise TF-IDF cosine. Brackets: 95% bootstrap intervals over templates. Model audit: share of a random sample (126/130, 117/126 and 119/125 items of the types we kept) that GPT-5.6 Terra judged correct, with Wilson 95% intervals.

|  | **Breast** | **Pancreas** | **Melanoma** |
|---|---|---|---|
| Cells | 92,237 | 187,593 | 86,970 |
| Cell clusters | 20 | 16 | 26 |
| Genes on the panel | 313 | 474 | 382 |
| Annotated regions | 7 | 20 | 6 |
| Templates | 1,058 | 1,002 | 1,018 |
| Released items | 2,500 | 2,500 | 2,500 |
| Distinct templates used | 851 | 899 | 916 |
| Distinct SQL shapes | 1,454 | 1,063 | 1,060 |
| Distinct ST_ functions | 51 | 47 | 47 |
| Queries with joins | 84.4 [82, 87] | 85.8 [84, 88] | 86.0 [84, 88] |
| Single-row answers | 63.2 [60, 67] | 66.9 [64, 70] | 66.4 [63, 69] |
| Mean question similarity | 0.062 [0.060, 0.064] | 0.059 [0.057, 0.061] | 0.058 [0.057, 0.061] |
| Agree with re-derivation | 75.8 [73, 78] | 85.2 [83, 87] | 85.1 [83, 87] |
| Model audit (%) | 96.9 [92, 99] | 92.9 [87, 96] | 95.2 [90, 98] |

### Conventions

Many biological questions can be ambiguous. Do “T cells” include both the CD4+ and the CD8+ clusters? Does the average expression of a gene count the cells in which it was not detected? For each dataset, a conventions document fixes these choices (Section F). The generator, critic, and the evaluation models all receive the schema, the conventions and the vocabulary (Figure 4). Because the conventions make each answer unique, we can score answers by execution.

### Phase 3 Template generation

We defined an 18-category query classification 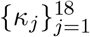. Each category *κ*_*j*_ uses its own subset of *ℱ*. Each template is a pair of a question and its `SQL` query, with slots for cell types, genes, spatial domains and distances. We wrote a few seed templates by hand, and an `LLM` (`Claude Haiku 4.5`) wrote more for each category from the schema, the vocabulary and the Phase 1 examples that use the category’s functions. Experimental biologists and pathology students refined the resulting 102 templates. To widen coverage, an `LLM` (`Claude Sonnet 5`) then wrote 1,127 more templates, given the existing templates and a profile of the data. We kept a new template only if its `SQL` ran on sampled slot values, returned a usable result, used one of the category’s functions, and passed the rule-based checks of Phase 5. This gives 1,229 templates for breast cancer (Figure 1 d). We later set aside three of the 18 categories (Phase 5), so the release covers 15. We ported these templates to the pancreas and melanoma sections by fixed rules (Section F).

### Phase 4 Initial NL2SQL pairs

An `LLM` (`Claude Sonnet 5`) instantiated each template, filling its slots with values from V and rewording the question. Generation ran in rounds, and templates with few accepted candidates received more, worded unlike those already accepted. Each candidate gets an `SQL` difficulty (easy, medium, hard or extra) from its joins, subqueries, groupings, set operations and aggregates, with the tier names of Spider (Yu et al., 2018), and a spatial difficulty (none, single, multi or advanced) from the number and kind of its `PostGIS` functions.

### Phase 5 Agentic verification

A near-duplicate filter first drops a candidate whose question, with numbers and entity names masked, has a token Jaccard similarity above 0.85 with an accepted question of the same template. Four independent checks then evaluate the surviving candidates (Figure 1). A semantic reviewer, 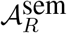, from another model family (GPT-5.6 Luna or Terra) writes its own `SQL` from the question, the schema and the conventions, without seeing the candidate. The two results must agree under the scorer of Section 5.3, except that either result may lack columns of the other. A syntactic validator, 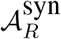, runs each candidate query twice and requires the same non-empty result both times. Rule-based checks flag common violations of the conventions, such as an average over sparse expression that drops zero counts, a `LEFT JOIN` undone by a filter on the joined table, a vague threshold word without a number, or a label that does not occur in the data. An agreeing pair must also agree on two half-sample copies of the database, each holding a fixed half of the cells. This test suite removes queries that agree on the full data by chance (Zhong et al., 2020). Finally, a model that does not see the question describes what the `SQL` computes, and a mismatch with the question is flagged (back-translation). Rows failing any check went to a judge agent (`Claude Opus 5` for breast cancer, `Claude Sonnet 5` for the other two datasets) that saw both queries and both results, and decided which query, if either, answers the question. A row entered the benchmark B only if: (*i*) 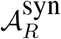 confirmed `SQL` execution, and (*ii*) the gold agreed with the re-derivation on the full and half-sample databases, or the judge accepted it and GPT-5.6 Terra, given the same evidence, made the same choice. Most released items agree with their re-derivation (Table 1).

### Selection, audits and splits

Blind audits by two model families removed 2–5% of items per dataset, and we set aside three categories that ask about geometric constructs rather than the tissue. GPT-5.6 Terra judged 95.0% of a random sample correct (95% CI 92.3–96.8). We split by groups of related templates and hold out the same groups in all three datasets (Section F). The released benchmark *ℬ*gold ⊆ *ℬ* (`SPaRTA`-Bench) holds 7,500 items: 33% are easy, 52% medium, 13% hard and 2% extra hard, and 31% use advanced spatial operations (Figure 2, Table 1).

**Figure 2.**
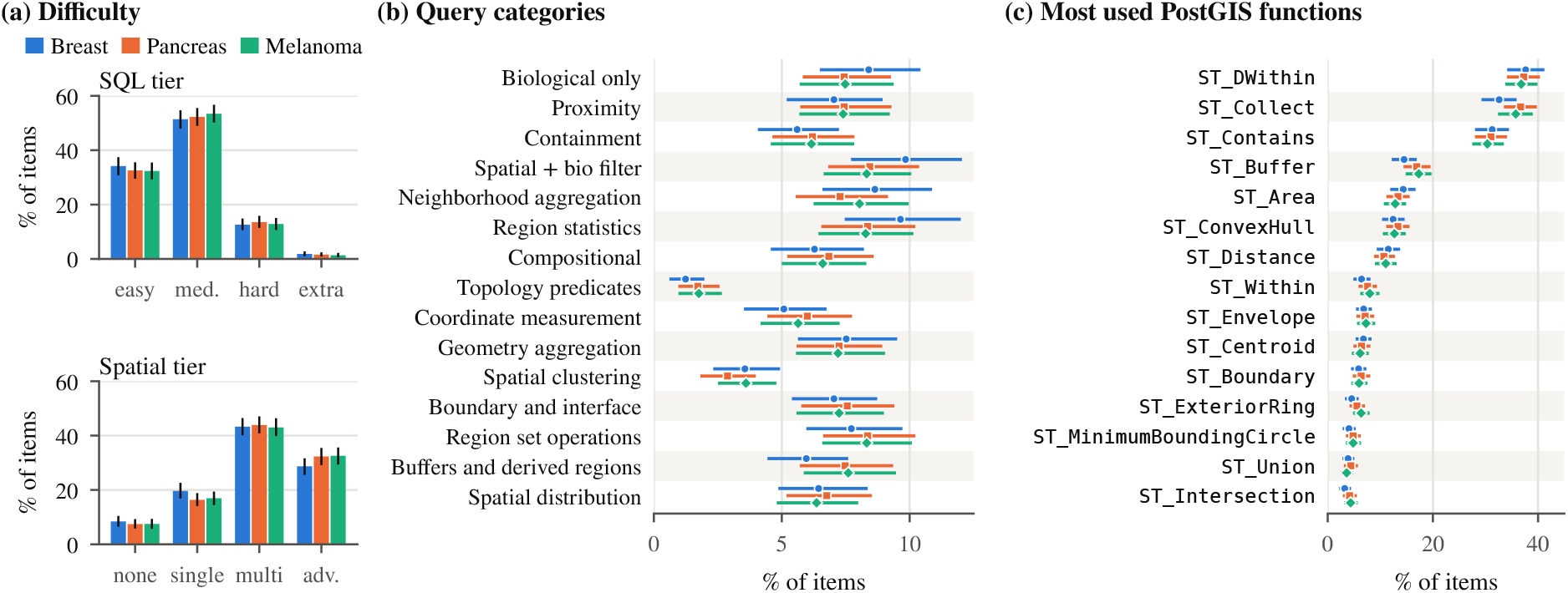
Characteristics of `SPaRTA`-Bench. (**a**) `SQL` and spatial difficulty levels. (**b**) The 15 query categories in each dataset. (**c**) Spatial functions in the released `SQL` queries. Shares of the full release, with 95% bootstrap intervals over templates.

### Human validation

Four annotators, three of them biology experts, label 720 items, two per item (Section F). They pass 89–92% of labels in each dataset (weighted by audit confidence for breast cancer), and both annotators fail 4 of 475 doubly labeled items. Raw agreement is 83% (Gwet’s AC1 0.81).

## 4 Failure Mode Analysis

Incorrect `SQL` queries fall into two broad categories: those that fail to execute and those that execute but return incorrect results. The latter are particularly concerning because execution alone cannot detect them. Among 6,555 incorrect predictions from six base models in direct mode, 41% failed to execute, 11% returned the correct rows but omitted a required column, and 48% executed but returned incorrect results.

By comparing these predictions with their gold queries (Section 4.1), we identified 15 failure modes across three categories (Table 8; examples in Section B). *Semantic* failures misinterpret biological or spatial terms, such as confusing “region 3” with spatial domain 3. *Faithfulness* failures alter the meaning of the question, for example by omitting a distance margin or reversing an expression threshold. *Structural* failures produce incorrect query logic, such as using an inner join that excludes cells with zero expression and consequently inflates the average.

Together, these 15 modes account for 79% of silent failures (95% CI: 74–83%). The benchmark verification pipeline (Section 3) is designed to catch such errors through explicit query conventions, rule-based checks, and execution on independent half-samples of the data.

### 4.1 How often models make each error

To count these errors at scale, we compare each failed prediction with its gold query on eleven decisions that the conventions fix, such as the clusters, genes or regions a term maps to, the spatial predicate and radius, the join under sparse expression storage and geometry construction. We parse both queries and flag each decision on which they differ. Correct predictions differ from the gold only by harmless rewrites, and the detectors flag at most 3.7% of them, so false alarms are rare.

The weaker base models fail first on term resolution (Table 2): Llama-3.1-8B and Mistral-7B map a biological term to the wrong labels in 45% and 46% of their failed answers, and Qwen3-8B in 33% when it answers without reasoning. Stronger models rarely make this error (4% for Gemma-4-12B) and fail instead on geometry construction and on spatial predicates and radii. The sparse-join error of Table 7 is frequent for every base model (12– 26% of failures). After fine-tuning, term errors fall to 7% and spatial predicate errors to 10%. For 54% of the remaining failures, none of the eleven decisions differs from the gold, so fine-tuning removes most of the errors that the conventions name.

**Table 2.**
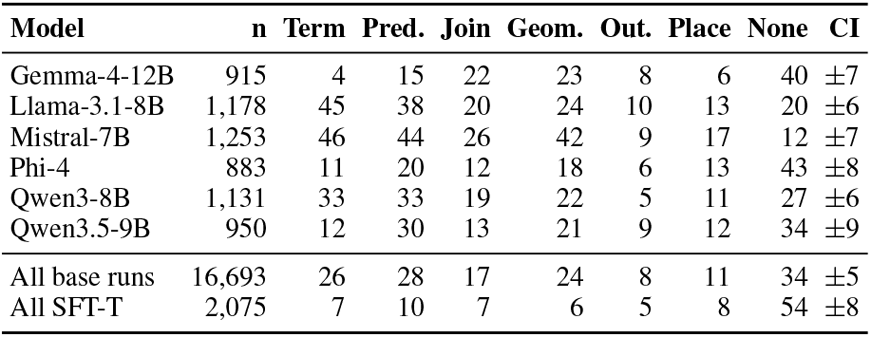
Share (%) of failed predictions that differ from the gold on each decision: term resolution, spatial predicate and radius, join under sparse storage, geometry construction, output shape and place-qualifier scope. None: no decision differs. CI: the largest half-width of a 95% bootstrap interval over template groups in the row. Model rows are base models in direct mode; “All base runs” pools the three modes of the six base models; “All SFT-T” pools the five fine-tuned models. A prediction can differ on several decisions. Each of the other five decisions appears in at most 4% of the failures of any row. The columns are defined in Table 14.

| **Model** | **n** | **Term** | **Pred.** | **Join** | **Geom.** | **Out.** | **Place** | **None** | **CI** |
|---|---|---|---|---|---|---|---|---|---|
| Gemma-4-12B | 915 | 4 | 15 | 22 | 23 | 8 | 6 | 40 | ±7 |
| Llama-3.1-8B | 1,178 | 45 | 38 | 20 | 24 | 10 | 13 | 20 | ±6 |
| Mistral-7B | 1,253 | 46 | 44 | 26 | 42 | 9 | 17 | 12 | ±7 |
| Phi-4 | 883 | 11 | 20 | 12 | 18 | 6 | 13 | 43 | ±8 |
| Qwen3-8B | 1,131 | 33 | 33 | 19 | 22 | 5 | 11 | 27 | ±6 |
| Qwen3.5-9B | 950 | 12 | 30 | 13 | 21 | 9 | 12 | 34 | ±9 |
| All base runs | 16,693 | 26 | 28 | 17 | 24 | 8 | 11 | 34 | ±5 |
| All SFT-T | 2,075 | 7 | 10 | 7 | 6 | 5 | 8 | 54 | ±8 |

## 5 Experiments

### 5.1 NL2SQL Generation using Different LLMs

We evaluate every model zero-shot, without few-shot examples or retrieval. Each question is an independent call that sees only the system prompt, which holds the schema, the conventions and the vocabulary (Figure 4), and the question. In *direct* mode the model writes one `SQL` query (reasoning off, greedy decoding). In *reasoning* (CoT) mode it reasons first. In *execution feedback* (agent) mode, we run the reasoning answer and return the error or the first rows, and the model may revise its query for up to three rounds (Section G).

### 5.2 Fine-tuning and domain adaptation

We fine-tune five of the six models with Low-Rank Adaptation (LoRA; Hu et al. 2022) on the training split of *ℬ*gold, with all three tissues in training (T) or with one tissue held out (D), and evaluate them on the test split in direct mode. The loss covers the target tokens only. We also train with DPO (Rafailov et al., 2023) on off-policy and on-policy pairs (Section G).

### 5.3 SPaRTA Evaluation Framework

Differences in row order, column names or extra columns change the result table without changing the answer, so a strict comparison underestimates a model. Partial credit errs the other way: a table with some correct rows can still support a wrong biological conclusion. We therefore use execution accuracy (Yu et al., 2018; Li et al., 2023), which scores each answer as right or wrong: EX = **1**[*R*(*T*_gold_) = *R*(*ϕ*(*T*_pred_))], where *R*(·) is the multiset of rows of a table and *ϕ* aligns the columns of the predicted table *T*_pred_ to those of the gold table *T*_gold_, ignoring row order. We accept extra columns if the gold columns line up row by row (*subset*), and a consistent renaming of the cluster ids that a clustering function assigns (*relabel*). A prediction must also match on both half-sample copies. We report 95% bootstrap intervals over template groups and compare runs with McNemar’s test for clustered pairs (Durkalski et al., 2003), with Holm’s correction within each family of comparisons. Section H.2 gives the column alignment and shows that model rankings within a setting are stable under exact scoring.

### 5.4 Results

On the 1,468 held-out test pairs of *ℬ*gold, open-weight models in direct mode reach 9.5–39.7% EX, with Phi-4 (39.7%) and Gemma-4-12B (37.5%) best (Table 3). Llama-3.1-8B and Mistral-7B write queries that fail to run on 54.5% and 57.4% of items, mostly through invalid syntax, columns absent from the schema or functions called on the wrong argument types, while Gemma-4-12B and Phi-4 fail to run on 11.2% and 14.5%. In the same mode, GPT-6.1-sol, `Claude` Sonnet 5.5 and Gemini 3.8 Flash reach 72.6%, 70.5% and 69.9% (mean of three runs). They score above the best open model zero-shot (Gemma-4-12B agent, 61.0%; Holm *p* ≤ 0.003) and do not differ from fine-tuned Gemma-4-12B (76.9%; Holm *p* ≥ 0.40).

**Table 3.** EX (%) on the 1,468 test items, pooled over the three datasets, with 95% bootstrap intervals over template groups in brackets (per dataset: Table 9). T: templates held out. D: templates and tissue held out. Significance uses the clustered McNemar test with Holm’s correction (*p <* 0.05). *: above the previous mode of the same model (CoT over direct, agent over CoT).↓: SFT-D below SFT-T, or DPO below SFT on the same split. Every SFT run beats its base model (*p <* 10^*−*6^). SFT-T over three seeds: Qwen3-8B 71.3 ± 1.8, Llama-3.1-8B 67.6 ± 3.6. Proprietary models answer in direct mode at each API’s low reasoning setting. †: earlier models of this family helped build the gold.

| **Model** | **Zero-shot**<br>**Direct** | **CoT** | **Agent** | **Fine-tuned, direct mode**<br>**SFT-T** | **SFT-D** | **DPO-T** | **DPO-D** |
|---|---|---|---|---|---|---|---|
| Gemma-4-12B | 37.5 [31, 44] | **58.9**<sup>\*</sup> [52, 65] | **61.0** [54, 68] | **76.9** [71, 82] | 71.3<sup>↓</sup> [66, 77] | 70.6<sup>↓</sup> [64, 76] | 64.0<sup>↓</sup> [58, 70] |
| Phi-4 | **39.7** [33, 46] | 40.4 [35, 46] | 46.3<sup>*</sup> [40, 52] | 74.8 [69, 80] | **73.6** [68, 79] | 67.6<sup>↓</sup> [61, 74] | 67.8<sup>↓</sup> [62, 73] |
| Qwen3.5-9B | 32.0 [26, 38] | 43.6<sup>*</sup> [38, 49] | 54.4<sup>*</sup> [49, 60] | – | – | – | – |
| Qwen3-8B | 22.5 [16, 29] | 52.9<sup>*</sup> [46, 59] | 57.9<sup>*</sup> [51, 65] | 73.3 [67, 79] | 70.6 [65, 76] | 70.8 [65, 76] | 63.9<sup>↓</sup> [57, 70] |
| on-policy DPO |  |  |  |  |  | 70.4 [63, 77] | 69.3 [63, 75] |
| Mistral-7B | 9.5 [5, 14] | 5.8 [3, 8] | 7.0 [4, 10] | 67.8 [61, 74] | 64.2 [57, 70] | 62.9 [56, 69] | 61.2 [54, 67] |
| Llama-3.1-8B | 12.2 [7, 18] | 11.6 [7, 16] | 18.1<sup>*</sup> [13, 23] | 63.5 [56, 70] | 63.2 [56, 70] | 58.1 [51, 64] | 58.2<sup>↓</sup> [51, 65] |
| *Proprietary reference, zero-shot, mean ± SD over three runs* |  |  |  |  |  |  |  |
| GPT-6.1-sol<sup>†</sup> | 72.6 ±0.3 [66, 79] | – | – | – | – | – | – |
| Claude Sonnet 5.5<sup>†</sup> | 70.5 ±0.4 [65, 76] | – | – | – | – | – | – |
| Gemini 3.8 Flash | 69.9 ±0.8 [63, 76] | – | – | – | – | – | – |

### Reasoning and feedback

Reasoning helps only the models trained to think: Qwen3-8B gains 30 points, Gemma-4-12B 21 and Qwen3.5-9B 12 (Table 3). Execution feedback adds 5 to 11 points more for four of the six models, mostly from queries that fail to run. It fixes 14% of first answers that fail to run, but only 4% of those that run and return a wrong result (Figure 3b). A model that sees a plausible table has no signal that it chose the wrong clusters or the wrong join.

**Figure 3.**
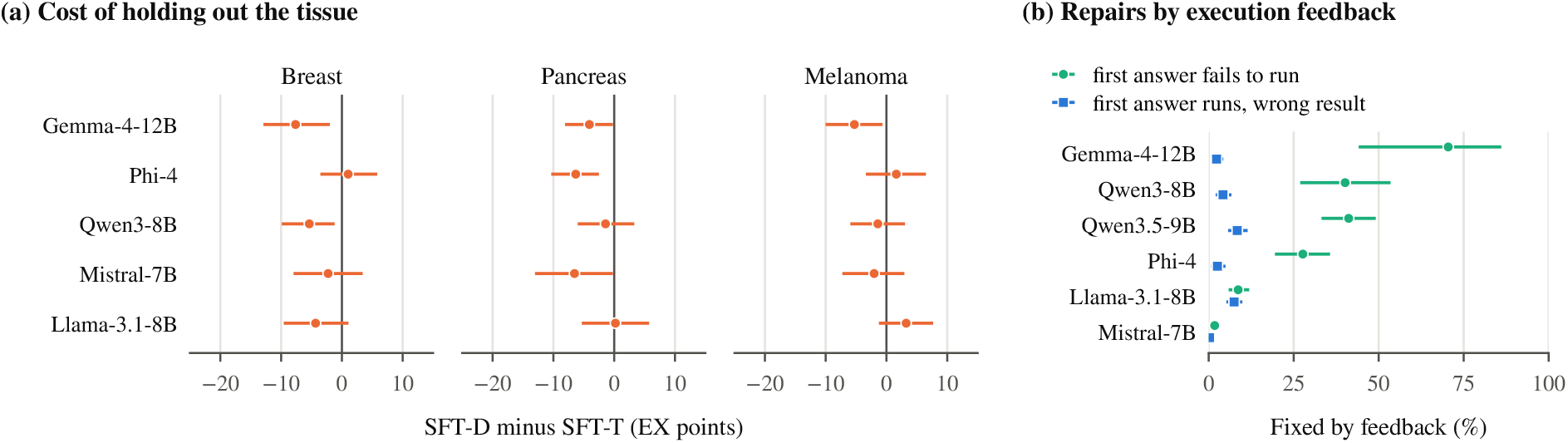
(**a**) Cost of holding out the tissue: SFT-D minus SFT-T on the same test items, per tissue. (**b**) Share of wrong first answers in agent mode that the final answer fixes, by the outcome of the first answer. Lines are 95% bootstrap intervals over template groups.

### Fine-tuning

SFT-T raises every model by 35 to 58 points, Mistral-7B the most (9.5% to 67.8%). Fine-tuned Gemma-4-12B beats its own zero-shot agent (76.9% against 61.0%, Holm *p <* 10^*−*4^) with one answer of about 110 tokens instead of reasoning and up to three rounds of feedback, and so does every fine-tuned model. The fine-tuned models differ little: Gemma-4-12B, Phi-4 and Qwen3-8B do not differ from one another (Holm *p* ≥ 0.43), and three seeds of Llama-3.1-8B span 7 points, so we treat gaps under 5 points as uncertain. Holding out the tissue (SFT-D) costs at most 5.7 points, and only Gemma-4-12B’s drop is significant (Holm *p* = 0.003). Breast and pancreatic cancer are harder to reach than melanoma (mean drops of 3.7, 3.6 and 0.8 points; Figure 3a). Off-policy DPO lowers EX for every model, by up to 7 points, significantly for six of ten runs, and on-policy pairs only recover SFT (Section G).

### By category

Fine-tuning helps most where a category needs one fixed construction that base models get wrong: spatial clustering rises from 7 to 85 and buffers from 37 to 91 (Table 4). Neighborhood aggregation, topology predicates, compositional queries and proximity stay near or below 50 after fine-tuning, and their failures turn on choices that the question leaves to the conventions (Section H).

**Table 4.** EX (%) by category on the test split, pooled over the three datasets. Dir. and Agent: base models in direct and agent mode, averaged over the six models. SFT-T: averaged over the five fine-tuned models. Gain is SFT-T minus base direct, over the five fine-tuned models. ±: half-width of a 95% bootstrap interval over the category’s 3–18 test templates.

| **Category** | **n** | **Dir.** | **Agent** | **SFT-T** | **Gain** |
|---|---|---|---|---|---|
| Biological only | 113 | 60.9 ±11 | 75.5 ±5 | 98.6 ±1 | +41.1 ±14 |
| Proximity | 107 | 15.6 ±10 | 27.7 ±14 | 50.5 ±21 | +37.4 ±23 |
| Containment | 85 | 16.9 ±15 | 31.0 ±18 | 68.9 ±18 | +54.6 ±19 |
| Spatial + bio filter | 134 | 11.2 ±8 | 22.6 ±12 | 63.6 ±17 | +53.0 ±16 |
| Neighborhood aggr. | 120 | 20.8 ±9 | 28.7 ±10 | 41.5 ±14 | +22.2 ±16 |
| Region statistics | 125 | 10.3 ±7 | 19.7 ±12 | 60.0 ±13 | +51.5 ±14 |
| Compositional | 98 | 8.5 ±9 | 23.8 ±13 | 45.7 ±14 | +37.6 ±14 |
| Topology predicates | 21 | 19.0 ±8 | 57.1 ±7 | 49.5 ±30 | +31.4 ±33 |
| Coordinate meas. | 82 | 35.2 ±13 | 49.0 ±12 | 89.5 ±8 | +57.8 ±12 |
| Geometry aggr. | 109 | 19.3 ±8 | 34.7 ±9 | 69.7 ±14 | +49.7 ±12 |
| Spatial clustering | 57 | 7.0 ±5 | 41.5 ±8 | 84.9 ±12 | +76.5 ±13 |
| Boundary/interface | 110 | 30.9 ±9 | 56.2 ±5 | 85.1 ±8 | +58.7 ±13 |
| Region set ops. | 103 | 21.7 ±9 | 51.0 ±10 | 73.4 ±14 | +52.6 ±13 |
| Buffers/derived | 104 | 37.2 ±12 | 42.1 ±16 | 90.8 ±7 | +52.9 ±12 |
| Spatial distribution | 100 | 61.7 ±10 | 72.5 ±5 | 95.6 ±3 | +32.4 ±10 |

## 6 Conclusion

We introduced `SPaRTA`, a benchmark of 7,500 verified NL2SQL pairs over three spatial-omics tissue sections. An agent-based pipeline generates the pairs, and each gold query must agree with an inde-pendent re-derivation, also on half-sample copies of the data, and pass blind audits. We identified 15 failure modes in which the query runs but returns a wrong answer. Execution feedback repairs queries that fail to run but rarely these silent failures. Fine-tuning raises five open-weight models to 63–77% execution accuracy with a single short answer, and the gains transfer to a tissue unseen in training. These results suggest a practical path toward privacy-preserving natural language interfaces to spatial-omics data, so that clinicians can query tissue maps as they query geospatial maps.

## 7 Limitations

### 7.1 Data scope limitation

The benchmark covers three spatial-omics sections, one each from breast cancer, pancreatic cancer and melanoma, all from one platform, `Xenium`. Other platforms differ in spatial resolution and gene panels. One sample per tissue limits the variety of cell types and expressed genes, and questions that span several samples, patients or time points are out of scope. Extending `SPaRTA` to other samples and conditions will require data from other technologies, tissue types and diseases.

### 7.2 Template specificity and semantic bias

All questions derive from 1,229 templates written for breast cancer, most of them by an `LLM`, and ported by rule to the other two tissues. We set aside question types that few biologists would ask, but this judgment was ours. Rewording and the nearduplicate filter make the questions varied, but they are not questions that clinicians asked. Questions that arise in clinical practice, such as multi-step reasoning across cell states, time-series comparisons or differential analysis across tissues, are absent because they do not fit the templates. Only sustained work with pathologists and clinicians can map the questions they ask.

### 7.3 Model-built gold

`LLM`s wrote the questions and the gold queries, and other `LLM`s checked and judged them. An item was kept only when models of two families agreed, which may favor the readings these models prefer. Model audits estimate that 93–97% of items are correct, and the second full audit did not run for pancreas. The human validation (Section 3) covers 720 items, and its breast cancer sample is not a simple random sample.

## 8 Ethical considerations

`SPaRTA` uses the publicly available GeoSQL-Bench for grounding the generative process, and three publicly available Xenium datasets, breast cancer (Janesick et al., 2023), pancreatic cancer (10x Genomics, 2023b) and melanoma (10x Genomics, 2023a), available from the 10x Genomics website. No patient-identifying information is involved. The generation and verification of the benchmark use proprietary models accessed with API keys (`Claude` Haiku 4.5, Sonnet 5 and Opus 5, GPT5.6 Luna and Terra). The fine-tuned models, however, are open-weight and run locally, so the approach can serve sensitive data without sending it to external services. The proprietary reference models receive only the public benchmark questions and schema. All experimental biologists involved in template selection produce and analyze spatial-omics datasets in their own research. Four annotators, three of them biology experts, validated the gold items (Section F). They labeled the items without seeing any model prediction and followed written instructions. A fine-tuned model can still return a wrong answer that looks plausible, so its outputs should be checked before they inform diagnosis or treatment. Finally, we used *ChatGPT* as a writing assistant for flow and grammar, and Claude (Anthropic) as an assistant for code and editing. All scientific decisions, interpretations, and core arguments were made by the authors.

## Acknowledgments

The authors would like to acknowledge the help of Dwaipayan Datta in the previous version of the project. Hirak Sarkar and Souvik Ghosh are partially supported by IRG-22-145-62 and the VICC GI SPORE (P50CA236733).

## A Template based Examples

We selected 6 template categories as a basis for generating `SQL` queries, and later extended them to 18, of which the release keeps 15 (Table 5). These templates span a wide range of spatial queries:

**Table 5.**
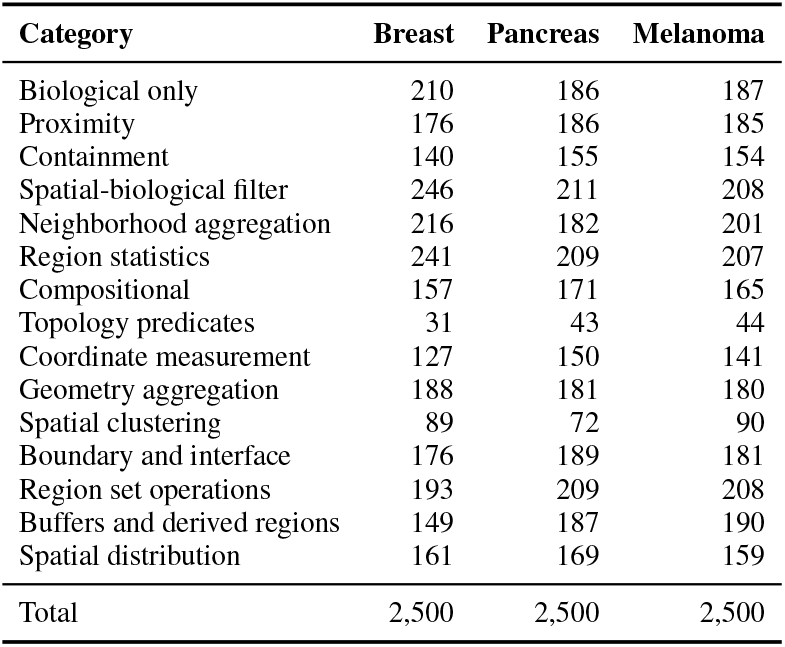
Released items per category. Proximity through Compositional are the six original categories. We set aside three categories of the 3,000-item sets: hulls and shape (200 items over the three datasets), tessellation and neighbor graph (440) and shape similarity (401) (Section 3).

| **Category** | **Breast** | **Pancreas** | **Melanoma** |
|---|---|---|---|
| Biological only | 210 | 186 | 187 |
| Proximity | 176 | 186 | 185 |
| Containment | 140 | 155 | 154 |
| Spatial-biological filter | 246 | 211 | 208 |
| Neighborhood aggregation | 216 | 182 | 201 |
| Region statistics | 241 | 209 | 207 |
| Compositional | 157 | 171 | 165 |
| Topology predicates | 31 | 43 | 44 |
| Coordinate measurement | 127 | 150 | 141 |
| Geometry aggregation | 188 | 181 | 180 |
| Spatial clustering | 89 | 72 | 90 |
| Boundary and interface | 176 | 189 | 181 |
| Region set operations | 193 | 209 | 208 |
| Buffers and derived regions | 149 | 187 | 190 |
| Spatial distribution | 161 | 169 | 159 |
| Total | 2,500 | 2,500 | 2,500 |

**Table 6.**
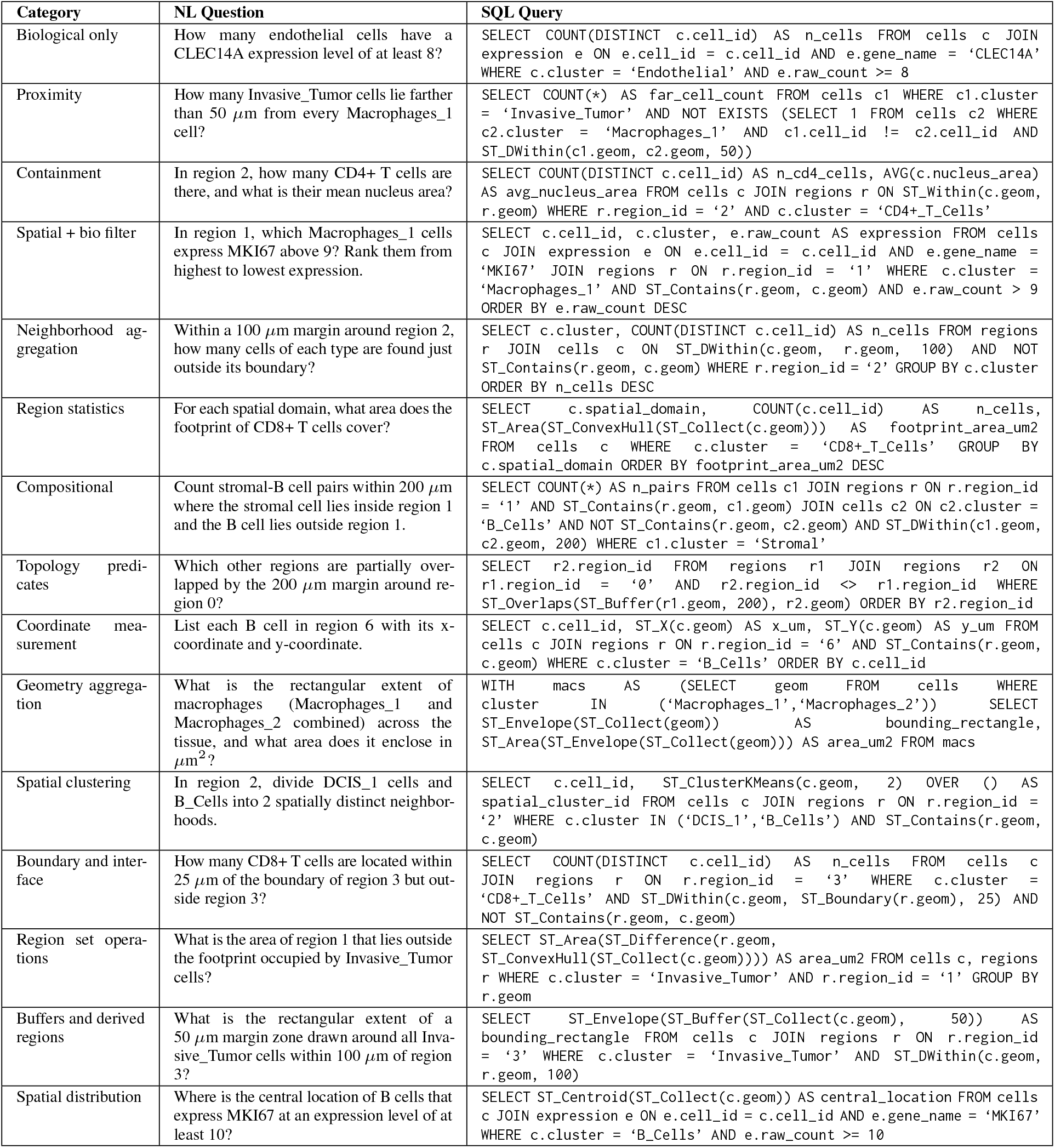
Released question–`SQL` pairs from the breast cancer section of `SPaRTA`-Bench, one for each of the 15 released query categories. Each gold query agrees with an independent re-derivation on the full data and on both half-sample copies.

| **Category** | **NL Question** | **SQL Query** |
|---|---|---|
| Biological only | How many endothelial cells have a CLEC14A expression level of at least 8? | SELECT COUNT(DISTINCT c.cell_id) AS n_cells FROM cells c JOIN expression e ON e.cell_id = c.cell_id AND e.gene_name = ‘CLEC14A’ WHERE c.cluster = ‘Endothelial’ AND e.raw_count >= 8 |
| Proximity | How many Invasive_Tumor cells lie farther than 50 µm from every Macrophages_1 cell? | SELECT COUNT(*) AS far_cell_count FROM cells c1 WHERE c1.cluster = ‘Invasive_Tumor’ AND NOT EXISTS (SELECT 1 FROM cells c2 WHERE c2.cluster = ‘Macrophages_1’ AND c1.cell_id != c2.cell_id AND ST_DWithin(c1.geom, c2.geom, 50)) |
| Containment | In region 2, how many CD4+ T cells are there, and what is their mean nucleus area? | SELECT COUNT(DISTINCT c.cell_id) AS n_cd4_cells, AVG(c.nucleus_area) AS avg_nucleus_area FROM cells c JOIN regions r ON ST_Within(c.geom, r.geom) WHERE r.region_id = ‘2’ AND c.cluster = ‘CD4+_T_Cells’ |
| Spatial + bio filter | In region 1, which Macrophages_1 cells express MKI67 above 9? Rank them from highest to lowest expression. | SELECT c.cell_id, c.cluster, e.raw_count AS expression FROM cells c JOIN expression e ON e.cell_id = c.cell_id AND e.gene_name = ‘MKI67’ JOIN regions r ON r.region_id = ‘1’ WHERE c.cluster = ‘Macrophages_1’ AND ST_Contains(r.geom, c.geom) AND e.raw_count > 9 ORDER BY e.raw_count DESC |
| Neighborhood ag-gregation | Within a 100 µm margin around region 2, how many cells of each type are found just outside its boundary? | SELECT c.cluster, COUNT(DISTINCT c.cell_id) AS n_cells FROM regions r JOIN cells c ON ST_DWithin(c.geom, r.geom, 100) AND NOT ST_Contains(r.geom, c.geom) WHERE r.region_id = ‘2’ GROUP BY c.cluster ORDER BY n_cells DESC |
| Region statistics | For each spatial domain, what area does the footprint of CD8+ T cells cover? | SELECT c.spatial_domain, COUNT(c.cell_id) AS n_cells, ST_Area(ST_ConvexHull(ST_Collect(c.geom))) AS footprint_area_um2 FROM cells c WHERE c.cluster = ‘CD8+_T_Cells’ GROUP BY c.spatial_domain ORDER BY footprint_area_um2 DESC |
| Compositional | Count stromal-B cell pairs within 200 µm where the stromal cell lies inside region 1 and the B cell lies outside region 1. | SELECT COUNT(*) AS n_pairs FROM cells c1 JOIN regions r ON r.region_id = ‘1’ AND ST_Contains(r.geom, c1.geom) JOIN cells c2 ON c2.cluster = ‘B_Cells’ AND NOT ST_Contains(r.geom, c2.geom) AND ST_DWithin(c1.geom, c2.geom, 200) WHERE c1.cluster = ‘Stromal’ |
| Topology predi-cates | Which other regions are partially over-lapped by the 200 µm margin around re-gion 0? | SELECT r2.region_id FROM regions r1 JOIN regions r2 ON r1.region_id = ‘0’ AND r2.region_id <> r1.region_id WHERE ST_Overlaps(ST_Buffer(r1.geom, 200), r2.geom) ORDER BY r2.region_id |
| Coordinate mea-surement | List each B cell in region 6 with its x-coordinate and y-coordinate. | SELECT c.cell_id, ST_X(c.geom) AS x_um, ST_Y(c.geom) AS y_um FROM cells c JOIN regions r ON r.region_id = ‘6’ AND ST_Contains(r.geom, c.geom) WHERE c.cluster = ‘B_Cells’ ORDER BY c.cell_id |
| Geometry aggrega-tion | What is the rectangular extent of macrophages (Macrophages_1 and Macrophages_2 combined) across the tissue, and what area does it enclose in µm<sup>2</sup>? | WITH macs AS (SELECT geom FROM cells WHERE cluster IN (‘Macrophages_1’,‘Macrophages_2’)) SELECT ST_Envelope(ST_Collect(geom)) AS bounding_rectangle, ST_Area(ST_Envelope(ST_Collect(geom))) AS area_um2 FROM macs |
| Spatial clustering | In region 2, divide DCIS_1 cells and B_Cells into 2 spatially distinct neighbor-hoods. | SELECT c.cell_id, ST_ClusterKMeans(c.geom, 2) OVER () AS spatial_cluster_id FROM cells c JOIN regions r ON r.region_id = ‘2’ WHERE c.cluster IN (‘DCIS_1’,‘B_Cells’) AND ST_Contains(r.geom, c.geom) |
| Boundary and inter-face | How many CD8+ T cells are located within 25 µm of the boundary of region 3 but out-side region 3? | SELECT COUNT(DISTINCT c.cell_id) AS n_cells FROM cells c JOIN regions r ON r.region_id = ‘3’ WHERE c.cluster = ‘CD8+_T_Cells’ AND ST_DWithin(c.geom, ST_Boundary(r.geom), 25) AND NOT ST_Contains(r.geom, c.geom) |
| Region set opera-tions | What is the area of region 1 that lies outside the footprint occupied by Invasive_Tumor cells? | SELECT ST_Area(ST_Difference(r.geom, ST_ConvexHull(ST_Collect(c.geom)))) AS area_um2 FROM cells c, regions r WHERE c.cluster = ‘Invasive_Tumor’ AND r.region_id = ‘1’ GROUP BY r.geom |
| Buffers and derived regions | What is the rectangular extent of a 50 µm margin zone drawn around all Inva-sive_Tumor cells within 100 µm of region 3? | SELECT ST_Envelope(ST_Buffer(ST_Collect(c.geom), 50)) AS bounding_rectangle FROM cells c JOIN regions r ON r.region_id = ‘3’ WHERE c.cluster = ‘Invasive_Tumor’ AND ST_DWithin(c.geom, r.geom, 100) |
| Spatial distribution | Where is the central location of B cells that express MKI67 at an expression level of at least 10? | SELECT ST_Centroid(ST_Collect(c.geom)) AS central_location FROM cells c JOIN expression e ON e.cell_id = c.cell_id AND e.gene_name = ‘MKI67’ WHERE c.cluster = ‘B_Cells’ AND e.raw_count >= 10 |

- *Proximity*: Spatial-omics data capture which cells lie close to each other. Genes that covary in space and cell types that border each other point to biological processes, such as gene–gene interactions or a tumor boundary that resists immune cells.
- *Containment*: Regions often signify localized activity in a tissue. Such queries reveal the cell-type composition of a biological compartment. The geometry of these regions can show how a disease changes tissue organization.
- *Spatial + Bio Filter*: Spatial patterns of gene expression shape disease progression and the tumor microenvironment. These queries threshold a gene’s expression and then select the spatial points that pass.
- *Neighborhood Aggregation*: Counts and averages over the neighbors of each cell, such as the number of CD8+ T cells within 100 *µ*m of a tumor cell, describe the local cellular environment of that cell.
- *Region Statistics*: Spatial density and autocorrelation (Moran’s I) help find spatially varying genes, groups of specific cell types and cellular niches.
- *Compositional*: A selection followed by a computation on the selected cells. The order of the `SQL` operations determines the result.

The other nine released categories cover questions without geometry (cell-type composition and expression), topology predicates between regions, coordinate measurement, geometry aggregation, spatial clustering (DBSCAN and k-means), tissue boundaries and interfaces, set operations on regions, buffers and derived regions, and spatial distribution. We also built hulls and shape descriptors, tessellation and neighbor graphs (Delaunay and Voronoi), and shape similarity (Hausdorff distance), and later set them aside (Section 3).

Together `SPaRTA`-Bench contains a rich set of NL2SQL pairs that covers a wide variety of spatialomics questions.

## B Complete NL2SQL Failure Modes in SPaRTA

We call a test prediction a *silent failure* when it runs without error and returns a result that differs from the gold. Of the 6,555 failed predictions of the six base models in direct mode, 2,676 (41%) do not run, 732 (11%) return the gold rows but lack a gold column, and 3,147 (48%) are silent. After finetuning (SFT-T), 68% of the 2,109 failures are silent, because fine-tuned models rarely write queries that fail to run.

We assign failure modes with the detectors of Section 4.1. Each mode is a difference from the gold on a decision that the conventions fix (Table 8). We leave out two of the eleven decisions. The denominator detector flags correct predictions almost as often as silent failures (2.0% against 2.6%), and whether cells without neighbors appear in the output is a convention rather than a wrong answer. The remaining 15 modes cover 79% of the silent failures of the base models (95% CI 74–83) and 48% of those of the SFT-T models (40–55). The rest differ from the gold in ways that the detectors do not name. Table 7 shows three silent failures in full.

**Table 7.**
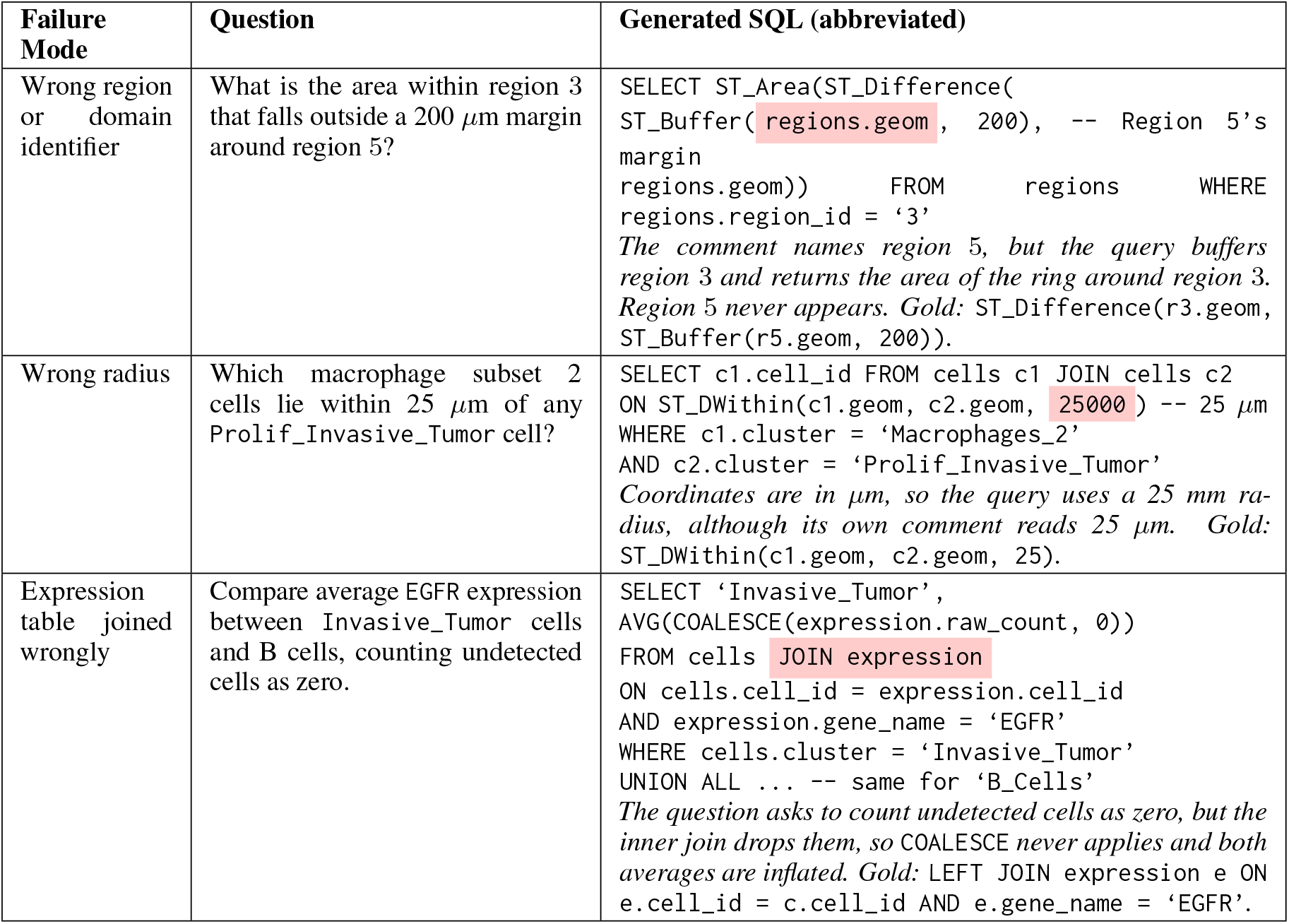
Three silent failures of base Mistral-7B in direct mode on the test split, one from each area. Each query runs and returns a plausible answer that differs from the gold, and the named mode is the only difference that the detectors find. Highlighted regions mark the erroneous fragment.

| **Failure Mode** | **Question** | **Generated SQL (abbreviated)** |
|---|---|---|
| Wrong region or domain identifier | What is the area within region 3 that falls outside a 200 µm margin around region 5? | SELECT ST_Area(ST_Difference(<br>ST_Buffer(regions.geom, 200), -- Region 5’s margin<br>regions.geom)) FROM regions WHERE regions.region_id = ‘3’<br>*The comment names region 5, but the query buffers region 3 and returns the area of the ring around region 3. Region 5 never appears. Gold:* ST_Difference(r3.geom, ST_Buffer(r5.geom, 200)). |
| Wrong radius | Which macrophage subset 2 cells lie within 25 µm of any Prolif_Invasive_Tumor cell? | SELECT c1.cell_id FROM cells c1 JOIN cells c2<br>ON ST_DWithin(c1.geom, c2.geom, 25000) -- 25 µm<br>WHERE c1.cluster = ‘Macrophages_2’<br>AND c2.cluster = ‘Prolif_Invasive_Tumor’<br>*Coordinates are in µm, so the query uses a 25 mm ra-dius, although its own comment reads 25 µm. Gold:* ST_DWithin(c1.geom, c2.geom, 25). |
| Expression table joined wrongly | Compare average EGFR expression between Invasive_Tumor cells and B cells, counting undetected cells as zero. | SELECT ‘Invasive_Tumor’,<br>AVG(COALESCE(expression.raw_count, 0))<br>FROM cells JOIN expression<br>ON cells.cell_id = expression.cell_id<br>AND expression.gene_name = ‘EGFR’<br>WHERE cells.cluster = ‘Invasive_Tumor’<br>UNION ALL ... -- same for ‘B_Cells’<br>*The question asks to count undetected cells as zero, but the inner join drops them, so* COALESCE *never applies and both averages are inflated. Gold:* LEFT JOIN expression e ON e.cell_id = c.cell_id AND e.gene_name = ‘EGFR’. |

Table 8 lists the 15 failure modes. Each row gives a query that a base model wrote in direct mode, in which the mode is the only difference from the gold that the detectors find, and the share of silent failures that show the mode.

**Table 8.**
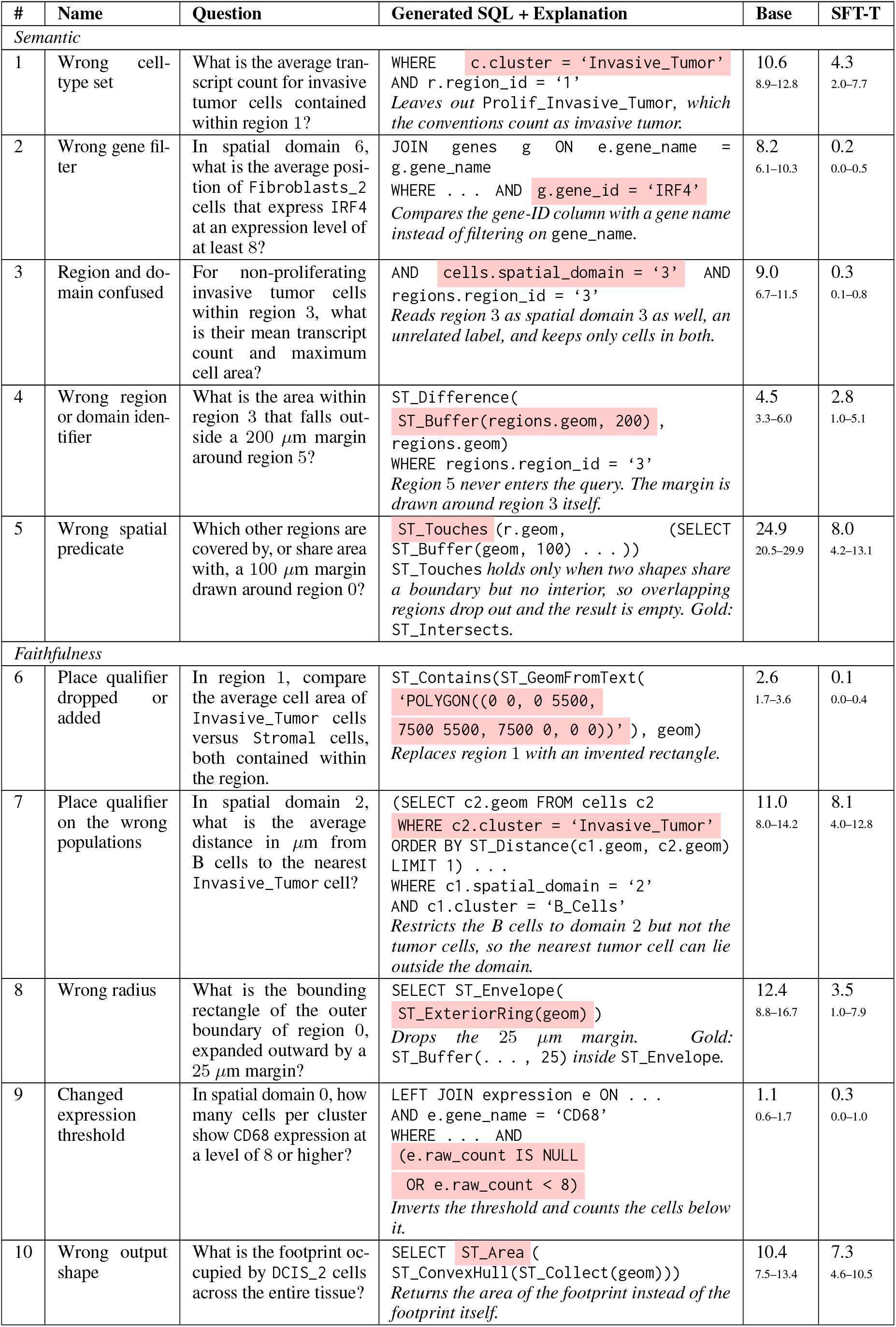
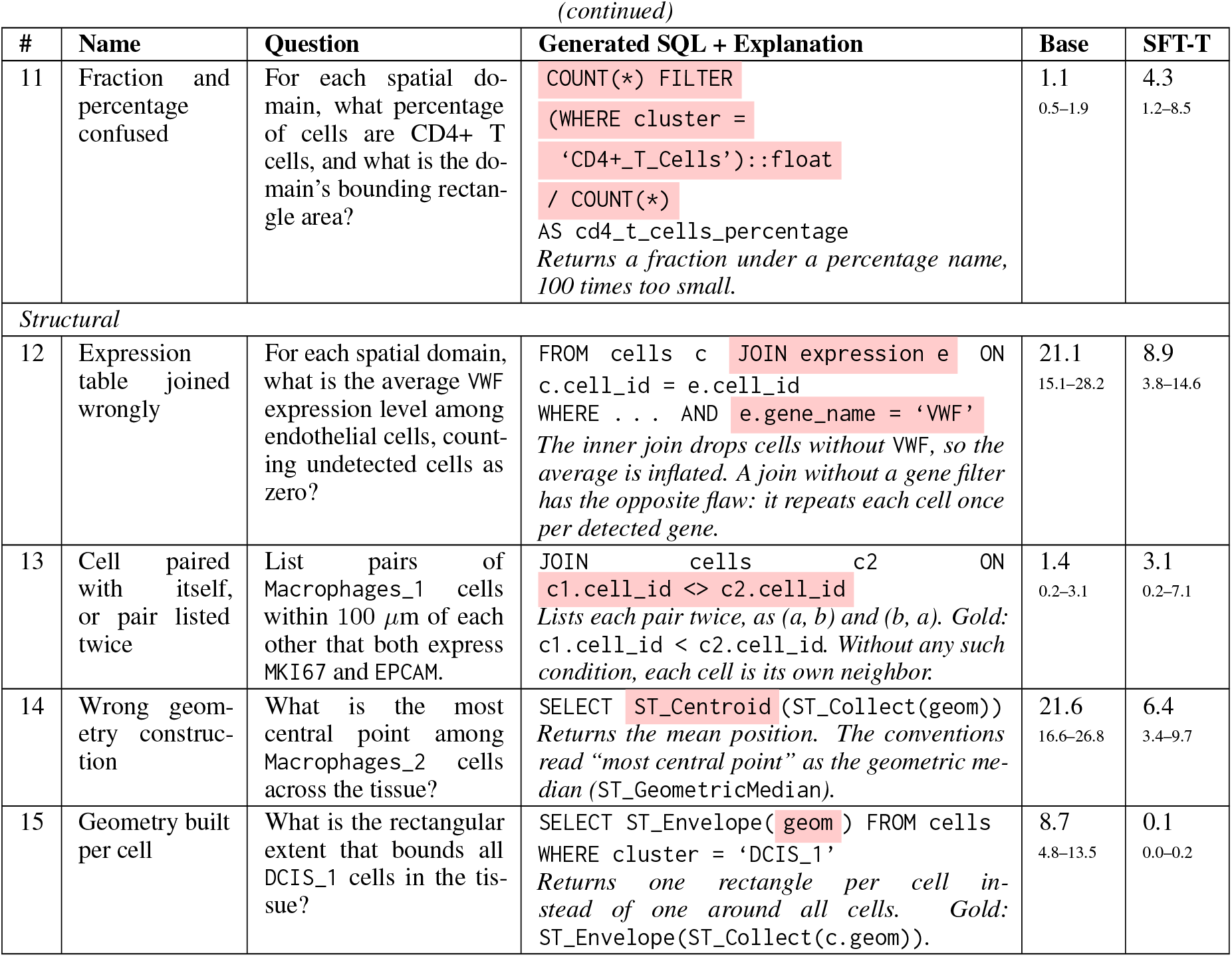
The 15 silent failure modes, measured on the test split. A silent failure runs and returns a result that differs from the gold. Each row shows a query that a base model wrote in direct mode (abbreviated; highlighted : the error), with a brief explanation in italics. Base and SFT-T: share (%) of silent failures that show the mode, for the six base models in direct mode (3,147 failures) and the five SFT-T models (1,433), with 95% bootstrap intervals over template groups. A failure can show several modes.

| **#** | **Name** | **Question** | **Generated SQL + Explanation** | **Base** | **SFT-T** |
|---|---|---|---|---|---|
| *Semantic* |  |  |  |  |  |
| 1 | Wrong cell-type set | What is the average tran-script count for invasive tumor cells contained within region 1? | WHERE c.cluster = ‘Invasive_Tumor’<br>AND r.region_id = ‘1’<br>*Leaves out* Prolif_Invasive_Tumor, *which the conventions count as invasive tumor.* | 10.6<br>8.9–12.8 | 4.3<br>2.0–7.7 |
| 2 | Wrong gene fil-ter | In spatial domain 6, what is the average posi-tion of Fibroblasts_2 cells that express IRF4 at an expression level of at least 8? | JOIN genes g ON e.gene_name = g.gene_name<br>WHERE . . . AND g.gene_id = ‘IRF4’<br>*Compares the gene-ID column with a gene name instead of filtering on* gene_name. | 8.2<br>6.1–10.3 | 0.2<br>0.0–0.5 |
| 3 | Region and do-main confused | For non-proliferating invasive tumor cells within region 3, what is their mean transcript count and maximum cell area? | AND cells.spatial_domain = ‘3’ AND regions.region_id = ‘3’<br>*Reads region 3 as spatial domain 3 as well, an unrelated label, and keeps only cells in both.* | 9.0<br>6.7–11.5 | 0.3<br>0.1–0.8 |
| 4 | Wrong region or domain iden-tifier | What is the area within region 3 that falls out-side a 200 µm margin around region 5? | ST_Difference(<br>ST_Buffer(regions.geom, 200) ,<br>regions.geom)<br>WHERE regions.region_id = ‘3’<br>*Region 5 never enters the query. The margin is drawn around region 3 itself.* | 4.5<br>3.3–6.0 | 2.8<br>1.0–5.1 |
| 5 | Wrong spatial predicate | Which other regions are covered by, or share area with, a 100 µm margin drawn around region 0? | ST_Touches (r.geom, (SELECT ST_Buffer(geom, 100) . . . ))<br>ST_Touches *holds only when two shapes share a boundary but no interior, so overlapping regions drop out and the result is empty. Gold:* ST_Intersects. | 24.9<br>20.5–29.9 | 8.0<br>4.2–13.1 |
| *Faithfulness* |  |  |  |  |  |
| 6 | Place qualifier dropped or added | In region 1, compare the average cell area of Invasive_Tumor cells versus Stromal cells, both contained within the region. | ST_Contains(ST_GeomFromText(<br>‘POLYGON((0 0, 0 5500,<br>7500 5500, 7500 0, 0 0))’ ), geom)<br>*Replaces region 1 with an invented rectangle.* | 2.6<br>1.7–3.6 | 0.1<br>0.0–0.4 |
| 7 | Place qualifier on the wrong populations | In spatial domain 2, what is the average distance in µm from B cells to the nearest Invasive_Tumor cell? | (SELECT c2.geom FROM cells c2<br>WHERE c2.cluster = ‘Invasive_Tumor’<br>ORDER BY ST_Distance(c1.geom, c2.geom)<br>LIMIT 1) . . .<br>WHERE c1.spatial_domain = ‘2’<br>AND c1.cluster = ‘B_Cells’<br>*Restricts the B cells to domain 2 but not the tumor cells, so the nearest tumor cell can lie outside the domain.* | 11.0<br>8.0–14.2 | 8.1<br>4.0–12.8 |
| 8 | Wrong radius | What is the bounding rectangle of the outer boundary of region 0, expanded outward by a 25 µm margin? | SELECT ST_Envelope(<br>ST_ExteriorRing(geom) )<br>*Drops the 25 µm margin. Gold:* ST_Buffer(. . . , 25) *inside* ST_Envelope. | 12.4<br>8.8–16.7 | 3.5<br>1.0–7.9 |
| 9 | Changed expression threshold | In spatial domain 0, how many cells per cluster show CD68 expression at a level of 8 or higher? | LEFT JOIN expression e ON . . .<br>AND e.gene_name = ‘CD68’<br>WHERE . . . AND<br>(e.raw_count IS NULL<br>OR e.raw_count < 8)<br>*Inverts the threshold and counts the cells below it.* | 1.1<br>0.6–1.7 | 0.3<br>0.0–1.0 |
| 10 | Wrong output shape | What is the footprint oc-cupied by DCIS_2 cells across the entire tissue? | SELECT ST_Area (<br>ST_ConvexHull(ST_Collect(geom)))<br>*Returns the area of the footprint instead of the footprint itself.* | 10.4<br>7.5–13.4 | 7.3<br>4.6–10.5 |
| **#** | **Name** | **Question** | **Generated SQL + Explanation** | **Base** | **SFT-T** |
| 11 | Fraction and percentage confused | For each spatial do-main, what percentage of cells are CD4+ T cells, and what is the do-main’s bounding rectan-gle area? | COUNT(\*) FILTER<br>(WHERE cluster =<br>‘CD4+_T_Cells’)::float<br>/ COUNT(\*)<br>AS cd4_t_cells_percentage<br>*Returns a fraction under a percentage name, 100 times too small.* | 1.1<br>0.5–1.9 | 4.3<br>1.2–8.5 |
| *Structural* |  |  |  |  |  |
| 12 | Expression table joined wrongly | For each spatial domain, what is the average VWF expression level among endothelial cells, count-ing undetected cells as zero? | FROM cells c JOIN expression e ON c.cell_id = e.cell_id<br>WHERE . . . AND e.gene_name = ‘VWF’<br>*The inner join drops cells without* VWF*, so the average is inflated. A join without a gene filter has the opposite flaw: it repeats each cell once per detected gene.* | 21.1<br>15.1–28.2 | 8.9<br>3.8–14.6 |
| 13 | Cell paired with itself, or pair listed twice | List pairs of Macrophages_1 cells within 100 µm of each other that both express MKI67 and EPCAM. | JOIN cells c2 ON c1.cell_id <> c2.cell_id<br>*Lists each pair twice, as (a, b) and (b, a). Gold:* c1.cell_id < c2.cell_id. *Without any such condition, each cell is its own neighbor.* | 1.4<br>0.2–3.1 | 3.1<br>0.2–7.1 |
| 14 | Wrong geom-etry construc-tion | What is the most central point among Macrophages_2 cells across the tissue? | SELECT ST_Centroid (ST_Collect(geom))<br>*Returns the mean position. The conventions read “most central point” as the geometric me-dian (*ST_GeometricMedian*).* | 21.6<br>16.6–26.8 | 6.4<br>3.4–9.7 |
| 15 | Geometry built per cell | What is the rectangular extent that bounds all DCIS_1 cells in the tis-sue? | SELECT ST_Envelope( geom ) FROM cells<br>WHERE cluster = ‘DCIS_1’<br>*Returns one rectangle per cell in-stead of one around all cells. Gold:* ST_Envelope(ST_Collect(c.geom)). | 8.7<br>4.8–13.5 | 0.1<br>0.0–0.2 |

## C Model Version Details

For our experiments, we use the following open-weight checkpoints from Hugging Face, served in bf16 with vLLM 0.27.1 on one NVIDIA RTX A6000 (48 GB) each, with each model’s own chat template.

- `Llama:` meta-llama/Llama-3.1-8B-Instruct
- `Mistral:` mistralai/Mistral-7B-Instruct-v0.3
- `Gemma:` google/gemma-4-12b-it
- `Qwen:` Qwen/Qwen3-8B and Qwen/Qwen3.5-9B
- `Phi:` microsoft/phi-4

For benchmark construction, we use the following proprietary models through an institutional API gateway between 24 September and 1 October 2026, and `Claude` Sonnet 5 also through the Anthropic Batch API on 1 October 2026 for part of the pancreas audit.

- `Claude:` claude-haiku-4-5-20251001 (original templates), claude-sonnet-5 and claude-opus-5
- `GPT:` gpt-5.6-luna and gpt-5.6-terra

### Proprietary reference

We ran GPT-6.1-sol (`gpt-6.1-sol`), `Claude` Sonnet 5.5 (`claude-sonnet-5-5`) and Gemini 3.8 Flash (`gemini-3.8-flash`) through each provider’s batch API. Each model receives the system prompt and direct-mode question of the open models (Figure 4) and may write up to 8,000 tokens, reasoning included. All three use the API’s low reasoning setting: adaptive thinking at effort `low` for `Claude`, reasoning effort `low` for GPT and thinking level `low` for Gemini. We send no temperature, so the three runs are independent samples, and we take the last `SQL` block of each reply, as for the open models. `Claude` Sonnet 5 and Opus 5 wrote and judged the gold, and GPT-5.6 Luna and Terra re-derived and audited it. No Gemini model took part.

**Figure 4.**
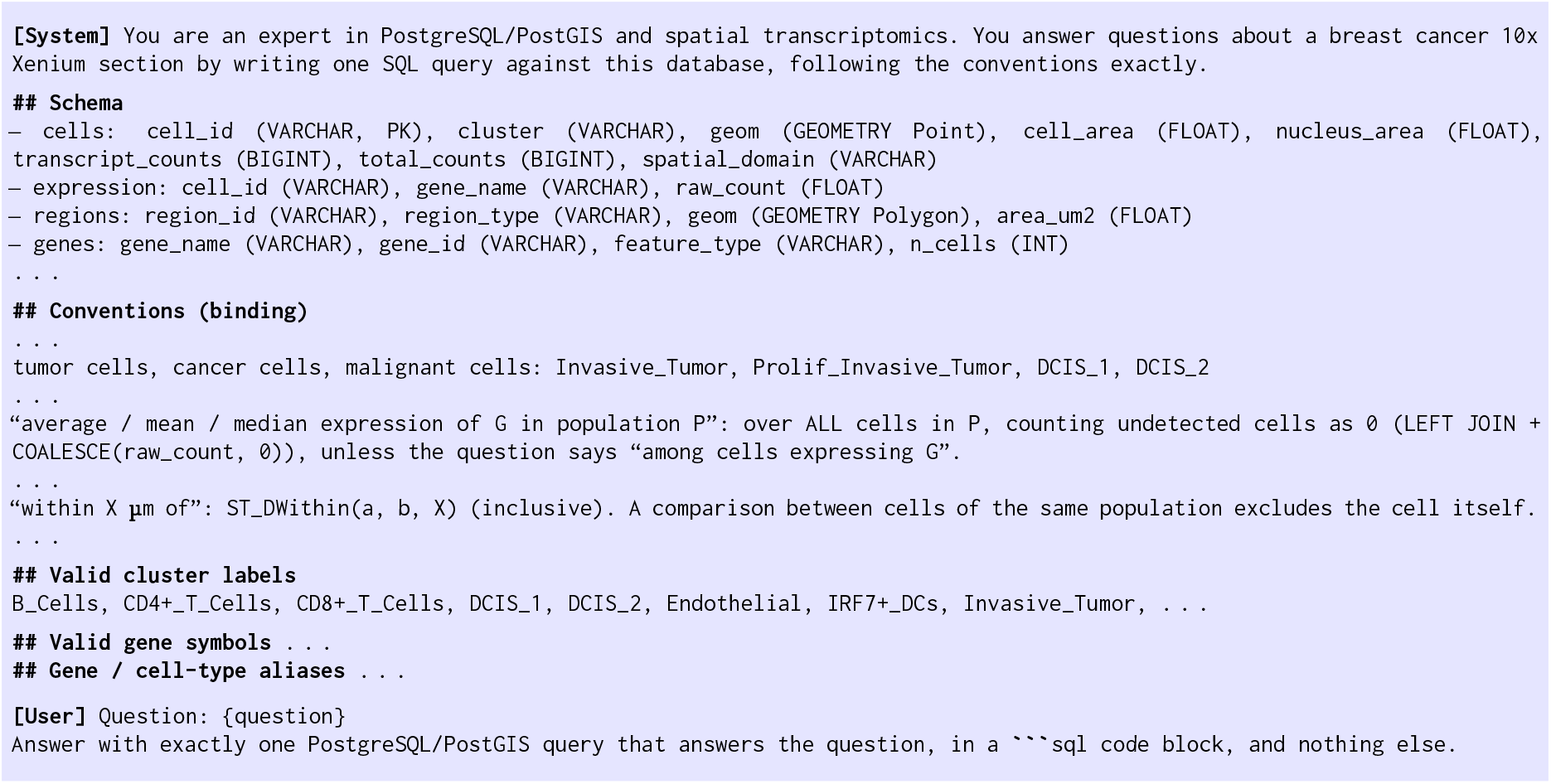
Excerpt of the system prompt for breast cancer (225 lines in full) and the direct-mode user message, used for all zero-shot and fine-tuning experiments. The schema grounds the model in concrete table definitions; the conventions fix the meaning of biological and spatial terms; the vocabulary lists every valid label.

## D Hyperparameter Details

All five models were fine-tuned using LoRA (Hu et al., 2022) on a single NVIDIA RTX A6000 (48 GB).

We apply LoRA with rank *r* = 16, *α* = 32, and dropout *p* = 0.05. For Llama-3.1-8B, Mistral-7B, Qwen3-8B, Phi-4 and Gemma-4-12B, adapters target all linear layers (query, key, value, output, gate, up, and down projections); for Gemma-4-12B, only those of the language model.

All models are trained for 2 epochs using our own trainer (PyTorch 2.13, Transformers 5.16, PEFT 0.20) with a cosine learning-rate schedule, peak learning rate 2×10^*−*4^, 3% warmup steps, and 48 examples per optimizer step. We measure the validation loss four times per epoch and keep the checkpoint with the lowest loss. All models used `bfloat16` precision and AdamW without weight decay, with gradients clipped at 1.0. Thinking models were trained with thinking mode disabled.

For DPO, both the initial policy and the reference model are the SFT adapter. We use *β* = 0.1 and an NLL weight of 1.0, train for 2 epochs with 10% warmup, and use a learning rate of 2×10^*−*5^ for off-policy pairs and 5×10^*−*6^ for on-policy pairs.

All examples of a dataset share one long system prompt (schema, conventions and vocabulary). We pack many examples after one copy of this prefix in a training row, with an attention mask that lets each example attend to the prefix and to itself only, and position ids that continue from the end of the prefix. Each example is then computed as if alone (logits agree within 10^*−*5^ in fp32) at a fraction of the cost. For models with sliding-window layers (Gemma 4) the mask also applies the window. This packing does not apply to linear-attention layers, so we do not fine-tune Qwen3.5-9B. Before evaluation we check that each served adapter reproduces the trainer’s validation loss on 32 examples.

## E Computing Environment

Benchmark construction (API calls and data processing) ran on a workstation with a 16-core AMD Ryzen Threadripper PRO 5955WX, 251 GB of RAM and PostgreSQL 18.4 with PostGIS.

All open model experiments are done in a research cluster consisting of four NVIDIA RTX A6000 (48 GB) GPU nodes. Training, evaluation and on-policy sampling took about 110 A6000 GPU-hours, one GPU per job. We used popular libraries including *PyTorch, HuggingFace Transformers, PEFT and vLLM*.

## F Benchmark Construction Details

### Spatial functions

The subset F of `PostGIS` functions (Figure 2c) centers on proximity (`ST_DWithin`, `ST_Distance`), containment (`ST_Contains`, `ST_Within`), region properties (`ST_Area`, `ST_Boundary`), spatial aggregation (`ST_Collect`, `ST_ClusterDBSCAN`, `ST_ConvexHull`) and derived regions (`ST_Buffer`, `ST_Union`, `ST_Difference`). The released queries use 54 distinct `ST_` functions. We restrict *ℱ* this way because the schema needs only points for cells and polygons for tissue regions. Other spatial operations such as routing, raster operations and coordinate transformations are less relevant to biological questions.

### Schema

The extraction produces two conceptually distinct types of spatial annotation. First, *spatial domain labels D* are per-cell categorical annotations (column `spatial_domain` in the `cells` table) encoding biologically meaningful groupings such as cell-type neighborhoods or niches. These are labels on cells, not geometric objects. Second, *regions* are spatially contiguous polygon geometries stored in a separate `regions` table, derived by computing polygons over cell coordinates per domain or loaded from an external `GeoJSON` annotation.

#### Conventions

The conventions document of each dataset fixes the clusters that each cell-type term covers, averages over all cells of a population with undetected cells counted as zero unless the question says otherwise, inclusive `ST_DWithin` at the distance the question states, no cell counted as its own neighbor, and the form of ratios, units and outputs (Figure 4).

### Categories

Six categories cover spatial proximity, containment, biological filtering, neighborhood aggregation, region-level statistics, and compositional multi-hop queries. The other nine released categories cover questions without geometry, topology predicates, coordinate measurement, geometry aggregation, spatial clustering, tissue boundaries and interfaces, region set operations, buffers and derived regions, and spatial distribution (Table 5).

#### Porting

We ported the breast cancer templates to the pancreas and melanoma sections with fixed rules: a cell type maps to the cluster with the same role, a gene stays if it is on the panel and is otherwise replaced by a marker with the same role, and regions map by size rank. Of the ported templates, 1,163 (pancreas) and 1,172 (melanoma) run on their data without errors, timeouts or empty results.

### Selection and wording

We kept at most three items per `SQL` shape (the same query with other values), dropped a question when its TF-IDF cosine similarity to a kept question of its category exceeded 0.8, and capped each category at 12% of a dataset. `Claude Sonnet 5` then rewrote stiff questions. A new wording was kept only if a rederivation from that wording alone returned the same answer.

### Final audits

The final audits saw the final wording and nothing of the pipeline. `Claude Sonnet 5` audited every item for correctness and naturalness and gave a confidence from 1 to 5. GPT-5.6 Luna also audited every breast cancer and melanoma item, and Terra broke the tie wherever Luna disputed an item that Sonnet passed. This second audit did not run for pancreas. The audits removed 54 of 3,230 breast cancer items, 114 of 3,300 pancreas items and 167 of 3,300 melanoma items. Most had wrong `SQL` (49, 76 and 118); the rest had ambiguous or unnatural questions. We also set aside items whose answer is a raw `PostGIS` record and items whose `SQL` repeats another item’s, and trimmed each dataset to 3,000 items by dropping the lowest-confidence item from the templates with the most items.

### Question types we set aside

Some templates ask about a geometric construct rather than the tissue: the edges of a Delaunay triangulation (“direct-neighbor links”), a cell’s Voronoi territory, a concave hull whose shape depends on a ratio the question cannot state (“tight outline”), or the Hausdorff distance between two footprints (“worst-case mismatch”). Few biologists would ask these questions, and their answers say more about the geometry than about the tissue. We removed the three categories built on them (hulls and shape, tessellation and neighborhood graph, shape similarity) and every other item whose gold query uses one of these constructs, 1,417 items in all. We then trimmed each dataset to 2,500 items with the rule above. The release holds 7,500 items in 15 categories, from 1,005 templates. We release the removed items with the reason for each removal. In a separate audit of 150 random items per dataset, drawn before this cut, 130, 126 and 125 items are of the types we kept. GPT-5.6 Terra judged 126, 117 and 119 of them correct (96.9%, 92.9% and 95.2%; 95.0% overall, Wilson 95% CI 92.3–96.8).

### Splits

Items made from the same template share their `SQL` structure, so a random split leaks it from training to test (Finegan-Dollak et al., 2018). We split by template instead. We first join templates that are variants of each other, share an `SQL` shape, or have questions with TF-IDF cosine of at least 0.85, which gives 902 template groups. We then assign groups to training (877 templates), validation (116) and test (236). The release uses 710, 97 and 198 of them. The same templates are held out in all three datasets, so the test set holds 485, 490 and 493 items (1,468 in all), and the training and validation sets hold 5,296 and 736. No test item shares a template, a coarse `SQL` shape, an exact query or an exact question with any item on the training side, including the candidate pool we use for preference pairs (Section 5.2). Removing the 13 test items whose question has a TF-IDF cosine of at least 0.9 with a training question changes no reported EX by more than 0.4 points. The median TF-IDF cosine between a test question and its nearest training question is 0.56, close to the 0.54 between training questions from different templates.

The template-held-out split (T) trains on the training templates of all three datasets. The dataset-held-out split (D) trains on the training templates of two datasets and tests on the test items of the third, so both the templates and the tissue (its cell types, genes and regions) are new at test time. T and D share the same test items, so their scores are directly comparable.

### Human validation

Four annotators label 240 items per dataset (720 in all). The items of a dataset form six blocks of 40, and each of the six annotator pairs labels one block, so every item has two independent labels and each annotator labels 360 items. An item passes when its question is clear and natural to a biologist and its `SQL` answers it under the conventions. Annotators may also answer *fail* or *maybe* (cannot decide without running more queries) and give a reason. The fourth annotator labeled again after the three experts explained how they handle ambiguous cases, and we report these labels. For all datasets the 240 items are a random sample. We drew the samples before the release cut, so pass rates count the 581 sampled items that are in the release. We report Cohen’s *κ* for each pair, and Krippendorff’s *α* and Gwet’s AC1 (Gwet, 2008) for each dataset, with 95% bootstrap intervals over items.

Annotators pass 89.4% of labels for pancreas (95% CI 85.7–93.0) and 88.9% for melanoma (85.4–92.2). For breast cancer the raw rate is 86.1% and the weighted rate is 91.7% (88.2–95.1). On the random block, one annotator passes 77% of items (61–90), and the same annotator passes 90% (82–97) of the reused items after weighting. The intervals overlap, but the weighted rate may overstate the rate of the release. Both annotators pass 84% of pancreas items and 81% of melanoma items, and most of the other items have one pass and one *maybe*. Both annotators fail 4 of the 475 doubly labeled release items. Over all 579 doubly labeled items, raw agreement on the three labels is 83%, *α* is 0.32 (0.22–0.42) and AC1 is 0.81 (0.77–0.85), with 82–85% in each dataset. Across the 14 pair– dataset cells labeled so far, *κ* ranges from −0.13 to 0.91 and AC1 from 0.62 to 0.97. Most items pass, and *κ* and *α* then stay low even when raw agreement is high (Feinstein and Cicchetti, 1990).

## G Inference and Training Data

### Direct

The model answers with one `SQL` query and nothing else. Reasoning is off and decoding is greedy. Fine-tuned models are trained for this mode.

### Reasoning (CoT)

The model reasons, then answers. Qwen3, Qwen3.5 and Gemma 4 use their native thinking mode; the other models are asked to reason in their reply. Decoding uses each model’s default sampling settings with a fixed seed per item.

### Execution feedback (agent)

After the reasoning answer, we run the query on the database (20second limit) and return the error or the first rows of the result. The model either keeps its answer or revises it, for up to three rounds. The final query is the last one the model wrote. The vocabulary in the prompt plays the role of a name-lookup tool, so execution is the agent’s only tool.

### Prompt construction

Each call consists of two parts: a system message containing the database schema *S*, the conventions, the vocabulary *V* and a fixed instruction, and a user message containing the natural language question *q* and the output instruction. The system message serves two roles: it grounds the model in the concrete table and column definitions and valid names (preventing hallucinated schema references), and it fixes the meaning of biological and spatial terms. The user message constrains the output format to one `SQL` query in a code block. An excerpt of the system prompt used across all models and all experiments is shown in Figure 4.

### Model-specific prompt handling

For openweight models, the system message is passed via the model’s native chat template. Most models accept a dedicated `system` role; otherwise, the schema is prepended to the first user turn. In direct mode, thinking models have chain-of-thought disabled at tokenization time to prevent reasoning traces from appearing in the output. All openweight models are run locally with vLLM (Kwon et al., 2023) in bf16 on one NVIDIA RTX A6000 GPU each, with greedy decoding (temperature 0) in direct mode.

### Training format

Each example is formatted using the model’s native chat template, with the direct-mode prompt of Section 5.1 (schema, conventions and vocabulary as a system prompt) and the gold `SQL` in a code block as the target. For models without a dedicated system role, the schema is prepended to the user turn. For thinking models, chain-of-thought is disabled at tokenization time.

We trained the adapters on the 3,000-item sets, before we set aside the question types of Section 3, and did not retrain them. The T split has 7,031 training examples: 6,361 items of those sets, 390 verified paraphrases of them and 280 items that passed the final audits but were trimmed. Of these, 1,059 (15%) belong to the types we later set aside. They share no template with the test items, and every test item belongs to the 2,500-item release. The validation set (875 examples) selects the check-point with the lowest loss. A D adapter trains on the same data without the held-out dataset. Appendix D gives the hyperparameters.

### Preference optimization

We start from each SFT adapter and train with DPO (Rafailov et al., 2023) plus a negative log-likelihood term on the chosen answer (Pang et al., 2024). The 2,086 off-policy pairs come from the benchmark pipeline: the chosen query is a verified gold, and the rejected query is one the pipeline found wrong, written by the generator or a checking model. We re-ran both queries to confirm that the rejected one fails or returns a different result, and drop all pairs from test templates. For Qwen3-8B we also build on-policy pairs: the SFT model answers each training and validation item four times at temperature 1, and a pair sets its most frequent correct query against its most frequent wrong one (1,717 training pairs for T).

### Seeds

To measure training noise, we retrain the T adapters of Qwen3-8B and Llama-3.1-8B with two more seeds.

## H Additional Results

### H.1 Main results in detail

#### Reasoning

Reasoning helps the models trained to think: Qwen3-8B gains 30 points, Gemma-4-12B 21 and Qwen3.5-9B 12. It does not help Phi-4, Llama or Mistral. For a given model and mode, scores on the three datasets differ by at most 10.2 points (Table 9).

**Table 9.** EX (%) per dataset on 485 (breast), 490 (pancreas) and 493 (melanoma) test items, with 95% bootstrap intervals over template groups. SFT-D on a dataset is the adapter trained on the other two. Proprietary rows: mean ± SD over three runs.

| Model | Run | Breast | Pancreas | Melanoma |
|---|---|---|---|---|
| Gemma-4-12B | direct | 40.0 [32, 48] | 38.8 [31, 46] | 33.9 [27, 41] |
|  | CoT | 61.6 [54, 69] | 56.9 [49, 65] | 58.0 [51, 65] |
|  | agent | 63.9 [56, 71] | 59.4 [51, 67] | 59.6 [52, 67] |
|  | SFT-T | 75.3 [68, 82] | 76.7 [70, 83] | 78.7 [73, 84] |
|  | SFT-D | 67.6 [61, 74] | 72.7 [66, 79] | 73.4 [67, 80] |
| Phi-4 | direct | 40.6 [33, 48] | 40.6 [34, 48] | 37.9 [31, 45] |
|  | CoT | 43.7 [36, 51] | 38.2 [32, 44] | 39.4 [33, 46] |
|  | agent | 49.1 [42, 56] | 44.9 [38, 52] | 45.0 [38, 52] |
|  | SFT-T | 73.2 [66, 80] | 77.3 [71, 83] | 73.8 [67, 80] |
|  | SFT-D | 74.2 [68, 80] | 71.0 [64, 77] | 75.5 [69, 82] |
| Qwen3.5-9B | direct | 37.7 [31, 44] | 27.6 [21, 34] | 30.8 [24, 37] |
|  | CoT | 44.9 [39, 51] | 45.3 [39, 51] | 40.6 [34, 47] |
|  | agent | 55.5 [49, 62] | 51.8 [45, 58] | 55.8 [49, 62] |
| Qwen3-8B | direct | 24.1 [17, 31] | 21.0 [14, 29] | 22.3 [16, 29] |
|  | CoT | 54.4 [47, 61] | 52.0 [45, 59] | 52.3 [45, 59] |
|  | agent | 60.6 [53, 68] | 56.7 [49, 64] | 56.4 [49, 64] |
|  | SFT-T | 71.3 [64, 78] | 73.1 [67, 79] | 75.5 [69, 82] |
|  | SFT-D | 66.0 [58, 73] | 71.6 [65, 78] | 74.0 [67, 80] |
| Llama-3.1-8B | direct | 14.4 [8, 22] | 12.0 [7, 18] | 10.1 [6, 15] |
|  | CoT | 14.8 [9, 20] | 9.8 [5, 15] | 10.3 [6, 14] |
|  | agent | 20.6 [14, 28] | 14.3 [10, 19] | 19.3 [14, 25] |
|  | SFT-T | 63.7 [56, 70] | 62.7 [55, 70] | 64.1 [57, 71] |
|  | SFT-D | 59.4 [52, 66] | 62.9 [54, 71] | 67.3 [60, 75] |
| Mistral-7B | direct | 10.1 [6, 15] | 10.0 [5, 16] | 8.5 [4, 13] |
|  | CoT | 4.9 [3, 7] | 6.9 [4, 10] | 5.5 [3, 8] |
|  | agent | 7.6 [4, 11] | 7.1 [4, 11] | 6.3 [3, 9] |
|  | SFT-T | 66.8 [59, 74] | 67.1 [60, 74] | 69.6 [62, 76] |
|  | SFT-D | 64.5 [57, 71] | 60.6 [52, 69] | 67.5 [60, 75] |
| GPT-6.1-sol | direct | 74.4 ±0.4 [67, 81] | 71.5 ±0.3 [64, 78] | 72.0 ±0.4 [65, 78] |
| Claude Sonnet 5.5 | direct | 73.5 ±0.7 [67, 79] | 68.8 ±0.3 [62, 75] | 69.3 ±0.8 [62, 76] |
| Gemini 3.8 Flash | direct | 73.3 ±0.8 [66, 80] | 68.8 ±0.4 [61, 76] | 67.6 ±1.2 [60, 74] |

#### Execution feedback

Execution feedback adds 1 to 11 points over reasoning, and Gemma-4-12B reaches 61.0%. The gain comes mostly from queries that fail to run. Over all six models, the final answer is right for 14% of first answers that raised an error (70% for Gemma, 40% for Qwen3, 2% for Mistral), but for only 4% of first answers that ran and returned a wrong result. By failure type, feedback fixes 4% of term resolution, predicate and sparse-join errors and 6% of geometry errors (Table 10). It rarely breaks a correct answer (53 of 3,164, 1.7%). A model that sees a plausible table has no signal that it chose the wrong clusters or the wrong join. Execution feedback, the standard tool-use remedy, does little against the silent failures of Section 4.

**Table 10.** Share (%) of wrong first answers in agent mode that the final answer fixes, with 95% bootstrap intervals over template groups and the number of such first answers in parentheses. First two rows: by outcome of the first answer. Last five rows: by the decision on which the first answer differs from the gold (Section 4.1); a first answer can differ on several.

| **First answer** | **Gemma** | **Llama** | **Mistral** | **Phi-4** | **Qwen3** | **Qwen3.5** | **All** |
|---|---|---|---|---|---|---|---|
| Does not run | 70 [44, 86] (44) | 9 [6, 12] (784) | 2 [0, 3] (966) | 28 [20, 36] (242) | 40 [27, 54] (152) | 41 [33, 49] (238) | 14 [12, 17] (2,426) |
| Runs, wrong result | 2 [1, 4] (559) | 7 [5, 10] (508) | 0 [0, 1] (405) | 3 [1, 5] (623) | 4 [2, 7] (546) | 8 [6, 12] (560) | 4 [3, 6] (3,201) |
| Term resolution | – (0) | 5 [3, 7] (571) | 1 [0, 1] (540) | 8 [0, 20] (48) | 14 [0, 35] (21) | 29 [13, 46] (28) | 4 [2, 5] (1,208) |
| Predicate and radius | 0 [0, 0] (51) | 6 [3, 9] (395) | 0 [0, 1] (548) | 6 [2, 12] (130) | 9 [3, 17] (67) | 20 [8, 36] (76) | 4 [3, 6] (1,267) |
| Join under sparse storage | 12 [0, 33] (17) | 5 [2, 9] (245) | 1 [0, 2] (299) | 4 [0, 9] (78) | 6 [0, 21] (35) | 14 [4, 27] (44) | 4 [2, 6] (718) |
| Geometry construction | 7 [0, 20] (61) | 5 [2, 9] (244) | 0 [0, 0] (475) | 13 [4, 23] (144) | 19 [7, 35] (90) | 14 [6, 24] (120) | 6 [3, 10] (1,134) |
| Output shape | 0 [0, 0] (36) | 7 [1, 14] (71) | 2 [0, 4] (102) | 10 [2, 19] (63) | 10 [0, 22] (29) | 15 [4, 28] (66) | 7 [4, 11] (367) |

#### Fine-tuning

SFT on the T split raises every model by 35 to 58 points, to 63.5–76.9%. Finetuned Qwen3-8B (73.3%) also beats its own agent (57.9%; Holm *p <* 10^*−*4^). Each fine-tuned model writes one answer of 89–124 output tokens on average. After fine-tuning, term errors, the main failure of the weaker models, fall from 26% to 7% of failures (Table 2).

#### Training noise

With one seed, 5 of the 10 pairwise differences between the SFT-T models are significant (Holm-adjusted *p <* 0.05). Gemma-4-12B, Phi-4 and Qwen3-8B do not differ from one another (*p* ≥ 0.43), and Mistral-7B differs from neither Qwen3-8B nor Llama-3.1-8B. Training noise adds to this. Three seeds of Qwen3-8B give 69.8–73.3% (s.d. 1.8), and three seeds of Llama-3.1-8B give 63.5, 69.1 and 70.1% (s.d. 3.6), with similar validation loss (0.110–0.115). Llama’s best seeds over-take Mistral-7B (67.8%, one seed). McNemar’s test accounts for the sampling of items, not for training noise. We therefore treat gaps of less than 5 points between fine-tuned models as uncertain.

#### A held-out tissue

Models trained on two tissues and tested on the third (SFT-D) lose at most 5.7 points against SFT-T, and only Gemma’s drop is significant (76.9 to 71.3, Holm-adjusted *p* = 0.003). Phi-4 and Llama lose at most 1.2 points. Breast and pancreatic cancer are harder to reach than melanoma. Four of five models lose on each of them, by up to 7.6 points on breast cancer (Gemma) and 6.5 on pancreatic cancer (Mistral) (Table 9). Most of what the models learn transfers to new cell types, genes and regions, which suggests they learn the conventions and the `SQL` rather than the vocabulary of one tissue.

#### Preference optimization

Off-policy DPO never improves over SFT. It lowers EX for every model on both splits, by up to 7.3 points, and six of ten drops are significant after correction (Holm-adjusted *p* ≤ 0.012). Execution errors rise (Phi-4 from 3.1% to 10.4% on T, Mistral from 6.1% to 12.7%). The rejected queries were written by other models, so the pairs push down text the policy would rarely produce, which is consistent with likelihood displacement in off-policy DPO (Razin et al., 2025). On-policy pairs remove the drop for Qwen3-8B on D: 69.3% against 63.9% with off-policy pairs (Holm-adjusted *p <* 0.001), and execution errors on D halve (4.6% against 9.1%). They do not differ from SFT (70.4% and 69.3% against 73.3% and 70.6%; Holm-adjusted *p* = 0.72).

#### Reasoning after fine-tuning

SFT on `SQL`-only targets removes the benefit of reasoning. With thinking on, fine-tuned Qwen3-8B scores 66.7%, below its own direct answers (73.3%; Holm *p* = 0.005). Fine-tuned Gemma-4-12B stops reasoning altogether: with thinking on it answers at once (75.9%). Combining the two gains would need training targets that keep the reasoning.

#### H.2 Metric validity

#### Column alignment

For `SPaRTA`, we first align columns via an injective mapping *ϕ* : *C*(*T*pred) → *C*(*T*gold). For each column *c*_*p*_, we assign *ϕ*(*c*_*p*_) = *c*_*g*_ when the multiset of values 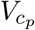 matches 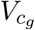, regardless of column names and order. The aligned columns must then also match row by row. Numbers match within a relative tolerance of 10^*−*3^ or the rounding of either side, labels match regardless of case and underscores, and geometries match within a Hausdorff distance of 0.01 *µ*m.

We accept two relaxations. A result with extra columns matches if the gold columns line up row by row (*subset*). A result whose group identifiers are a consistent renaming of the gold’s matches when the identifiers are cluster ids that a clustering function assigns arbitrarily, as with k-means and DBSCAN (*relabel*). A prediction that matches on the full database must also match on both halfsample copies (Section 3), so a query that is right only by chance counts as wrong. Gold and prediction each run with a 120-second limit and 4 GB of memory. A prediction that exceeds either limit counts as wrong (at most 0.9% of the predictions of any run).

We report 95% confidence intervals from a bootstrap over template groups, since items from one template are not independent. We compare two runs on the same items with McNemar’s test for clustered pairs (Durkalski et al., 2003), with template groups as clusters, and correct for multiple comparisons with Holm’s method within each family of comparisons that supports one claim.

#### Metric validity

We also rescore every run with no relaxations (*exact*; Table 11). The ranking of models within a setting is stable: Kendall’s *τ* against the EX ranking is 1.0 in five of seven settings and 0.8 in the other two.

**Table 11.** EX (%) with and without relaxations, with 95% bootstrap intervals over template groups. Exact: no relaxations. Proprietary rows are means over runs.

| Model | Run | Exact | EX |
|---|---|---|---|
| Gemma-4-12B | CoT | 58.6 [51, 65] | 58.9 [52, 65] |
|  | agent | 60.7 [53, 67] | 61.0 [54, 68] |
|  | SFT-T | 72.3 [66, 78] | 76.9 [71, 82] |
| Phi-4 | agent | 45.2 [39, 51] | 46.3 [40, 52] |
|  | SFT-T | 68.9 [62, 75] | 74.8 [69, 80] |
| Qwen3-8B | CoT | 51.5 [45, 58] | 52.9 [46, 59] |
|  | agent | 56.7 [50, 63] | 57.9 [51, 65] |
|  | SFT-T | 63.9 [57, 71] | 73.3 [67, 79] |
| Qwen3.5-9B | agent | 50.2 [44, 56] | 54.4 [49, 60] |
| GPT-6.1-sol | direct | 70.3 [63, 76] | 72.6 [66, 79] |
| Claude Sonnet 5.5 | direct | 59.7 [53, 66] | 70.5 [65, 76] |
| Gemini 3.8 Flash | direct | 69.5 [62, 76] | 69.9 [63, 76] |
| Mistral-7B | SFT-T | 65.5 [58, 72] | 67.8 [61, 74] |
| Llama-3.1-8B | SFT-T | 56.7 [49, 64] | 63.5 [56, 70] |

### H.3 Categories

#### Results by category

Table 4 breaks EX down by category. Fine-tuning helps most on categories whose queries follow one fixed construction that base models get wrong. Spatial clustering rises from 7.0 to 84.9 and buffers from 37.2 to 90.8. In base direct mode, 62% of clustering answers fail to execute; after fine-tuning, under 1% do. Four categories stay near or below 50 after fine-tuning: neighborhood aggregation (41.5), compositional queries (45.7), topology predicates (49.5) and proximity (50.5). Their remaining failures are mostly wrong results: 51.2% of neighborhood items return a wrong result, and 7.3% fail to execute. These failures turn on choices the question leaves to the conventions. In failed neighborhood predictions, the most frequent difference from the gold query is whether cells with no neighbor count (20%). In failed compositional predictions, it is which population a place qualifier restricts (27%).

A category’s score rests on 10 to 18 test templates (3 for topology), and templates within a category differ widely (Table 12). Under fine-tuning, the standard deviation of template EX ranges from 2 points (biological only) to 41 (proximity). Category differences of a few points are therefore within template noise. Topology predicates have 21 test items from 3 templates, too few to rank.

**Table 12.** Templates in the test split by category. Median and SD: template EX under SFT-T (mean over the five fine-tuned models). All: every fine-tuned model answers every item of the template. None: no base model answers any item of the template in direct mode. Brackets: 95% bootstrap intervals over the templates of the category.

| **Category** | **Tmpl.** | **Median** | **SD** | **All** | **None** |
|---|---|---|---|---|---|
| Biological only | 15 | 100.0 [98, 100] | 2.1 [1, 3] | 67 [40, 87] | 0 [0, 0] |
| Proximity | 14 | 41.3 [7, 89] | 40.9 [31, 45] | 14 [0, 36] | 43 [21, 71] |
| Containment | 12 | 84.8 [47, 98] | 28.2 [15, 33] | 25 [0, 50] | 42 [17, 67] |
| Spatial + bio filter | 18 | 84.7 [40, 92] | 34.5 [25, 40] | 22 [6, 44] | 50 [28, 72] |
| Neighborhood aggr. | 16 | 38.0 [18, 62] | 27.6 [20, 32] | 0 [0, 0] | 25 [6, 44] |
| Region statistics | 15 | 66.7 [38, 80] | 26.5 [18, 33] | 7 [0, 20] | 47 [20, 73] |
| Compositional | 12 | 43.2 [17, 68] | 26.6 [18, 32] | 0 [0, 0] | 42 [17, 67] |
| Topology predicates | 3 | 42.9 [31, 92] | 26.4 [0, 29] | 0 [0, 0] | 0 [0, 0] |
| Coordinate meas. | 11 | 100.0 [80, 100] | 15.4 [6, 20] | 55 [27, 82] | 9 [0, 27] |
| Geometry aggr. | 15 | 80.0 [56, 95] | 29.1 [16, 37] | 7 [0, 20] | 7 [0, 20] |
| Spatial clustering | 10 | 93.3 [68, 100] | 19.4 [9, 24] | 50 [20, 80] | 50 [20, 80] |
| Boundary/interface | 14 | 83.3 [73, 100] | 16.6 [10, 21] | 43 [21, 64] | 14 [0, 36] |
| Region set ops. | 13 | 85.0 [49, 94] | 25.7 [17, 30] | 8 [0, 23] | 8 [0, 23] |
| Buffers/derived | 17 | 100.0 [85, 100] | 20.9 [8, 30] | 59 [35, 82] | 0 [0, 0] |
| Spatial distribution | 13 | 98.0 [93, 100] | 5.0 [3, 6] | 46 [23, 69] | 0 [0, 0] |

#### Results by category in detail

Table 12 shows how each category’s score spreads over its templates. A template’s EX is the mean over its test items and the fine-tuned models.

#### Categories that fine-tuning solves

Biologicalonly questions, spatial distribution, buffers, coordinate measurement and spatial clustering all reach 84.9 or more after fine-tuning, and their median template scores 93 or more. Base models fail them for different reasons. In clustering, 212 of the 318 failed base answers do not execute, and 88 of those use `ST_ClusterKMeans` or `ST_ClusterDBSCAN` as an aggregate, though `PostGIS` defines both as window functions. No base model solves 50% of the clustering templates in direct mode. In buffers, most base answers run but construct the margin differently from the gold query (191 of 392 failures) or use another distance (132). Boundary and interface queries (85.1) follow the same course with a few hard templates left, such as the segment of a region’s edge near a cell type.

#### Preference tuning by category

Off-policy DPO lowers overall EX (Section 5.4), but not evenly. Against SFT-T, DPO-T loses 15.2 points on geometry aggregation, 13.3 on region statistics and 11.4 on topology predicates, and gains at most 2.1 (spatial clustering).

### H.4 Templates

#### Template difficulty

The 1,468 test items come from 198 templates (Table 13). Fine-tuning makes models agree on which templates are hard: the Spearman correlation of template EX between two models rises from 0.52 for base models in direct mode to 0.70 after SFT-T. Under SFT-T, 28% of templates are solved by every model and 3% by none, and the median template scores 84.4. A template that is hard in one tissue tends to be hard in the others (Spearman 0.76–0.82 between datasets), and 5–7% of templates differ by 50 points or more between two tissues. After fine-tuning, the `SQL` tier orders items by difficulty (82.1, 70.7, 53.8 and 31.7 from easy to extra hard), but the spatial tier does not: items that need one spatial operation score 57.9 and items that need an advanced one 72.2. Items with a subquery score 45.3 and items with three or more joins 48.2.

**Table 13.**
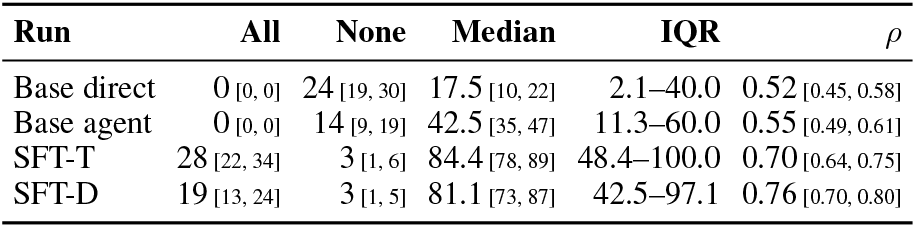
Template EX over the 198 test templates, for the five models we fine-tune. All (None): every (no) model of the run answers every (any) item of the template. Median and IQR: template EX averaged over the models of the run. *ρ*: Spearman correlation of template EX between two models of the run, averaged over pairs. Brackets: 95% bootstrap intervals over templates.

**Table 14.** Primary codes for failed predictions. Coders apply the rules in order: BENCHMARK, then CORRECT, then INVALID, then the first wrong decision in the order of the query’s logic (TERM, SPATIAL, GEOMETRY, AGGREGATION, OUTPUT). A second, independent error that would change the result by itself is a secondary code. Coders also rate their confidence from 1 to 3.

| **Code** | **Use it when** |
|---|---|
| TERM | A term is mapped to the wrong label or column: the cell type, the gene, the region or domain, or a place qualifier. |
| SPATIAL | The spatial relation is wrong: the predicate, the radius or its unit, whether neighbors on the boundary count, or whether a cell counts as its own neighbor. |
| GEOMETRY | A derived shape is built wrong (hull, buffer, envelope, centroid, union, boundary), or the wrong points go into it. |
| AGGREGATION | Rows are combined wrong: the join with sparse expression, grouping, a ratio’s denominator, distinct counting, or an expression threshold. |
| OUTPUT | The content is right but the form is wrong: missing or extra columns, ordering or limit, units, rounding. |
| INVALID | The query does not run. |
| BENCHMARK | The gold is wrong, or the question is ambiguous and the prediction is a defensible reading under the conventions. |
| CORRECT | The gold is right and the prediction also answers the question; the scorer was too strict. |
| OTHER | None of the above. |

#### Results by template

Table 13 summarizes template EX (the share of a template’s test items a model answers, pooled over datasets) for four runs. Base models disagree on which templates are hard. After fine-tuning, models agree more, and SFTT and SFT-D agree with each other (Spearman 0.90 between their template means). Fine-tuning raises 94% of test templates over base direct mode. Against the base agents, 11% of templates score lower under SFT-T, and 5 templates score more than 20 points lower, 2 of them in proximity and 2 in region set operations. Holding out the tissue costs 2.6 points per template on average. 8% of templates lose 20 points or more, and 2% gain 20 or more.

## I Schema Prompt

